# Progressive Vulnerability of Cortical Synapses in α-Synucleinopathy

**DOI:** 10.1101/2024.06.20.599774

**Authors:** Saroj Sah, Andrew D. Sauerbeck, Jyoti Gupta, Dayana Pérez-Acuña, Jacob E. Reiber, Dreson L. Russell, Vijay Singh, Lamisa Musarat, Laura A. Volpicelli-Daley, Michael J. Higley, Terrance T. Kummer, Thomas Biederer

## Abstract

α-Synuclein aggregates characterize α-synucleinopathies, including Parkinson’s disease and Dementia with Lewy bodies. A majority of people with these disorders experience cognitive decline, and its extent correlates with cortical α-synuclein pathology. Mechanisms by which pathology targets cortical circuits remain to be understood so that the debilitating non-motor impairments can be addressed. Overt neuronal loss is not a major feature of cortical pathology, but a reduction of presynaptic sites has been reported at late stages of PD. We here define excitatory synapses as neuronal loci affected by α-synuclein aggregation, showing that they are progressively lost, and identify temporal and spatial patterns of synaptic vulnerability. Results were obtained in a mouse model using intrastriatal injection of pre-formed α-synuclein fibrils to template the aggregation of endogenous α-synuclein in cortical neurons. Lewy neurite-like aggregates were predominantly observed in axons. Super-resolved imaging showed α-synuclein aggregation within cortical synapses and revealed that synaptic aggregation is most severe proximal to Lewy neurite-like structures and linked to the earliest detectable loss of excitatory synapses. Excitatory synapses also exhibited ultrastructural aberrations, including a redistribution of pre- and post-synaptic protein clusters away from contact sites and reduced synaptic vesicle size. As pathology advanced, VGLUT1-positive intracortical synapses, enriched in α-synuclein, were progressively vulnerable in this striatal seeding model, while VGLUT2-positive long-range synapses with minimal α-synuclein were spared. Inhibitory synapses were not affected. In agreement with a disruptive role of synaptic α-synuclein aggregation, super-resolved mesoscale imaging determined that synaptic but not non-synaptic α-synuclein pathology is correlated with excitatory synapse loss. Physiological recordings confirmed impaired excitatory neurotransmission. Pathology propagation tracked cortical connectivity, with intra-column synapse loss correlated between interconnected layers V and II/III. Contralateral areas receiving projections from pathologic layer V also exhibited synapse loss. These findings reveal synapses as principal cellular loci of α-synuclein pathology and define how molecular and circuit-based mechanisms underlie the cortical progression of synaptic pathology in α-synucleinopathies.

## Introduction

Intracellular protein aggregates of α-synuclein (α-syn), appearing as Lewy neurites and Lewy bodies, are a hallmark of synucleinopathies. This pathology is prominent in the cortex during the later stages of Parkinson’s disease (PD).^1–3^ Cognitive deficits involving the cortex are a prominent feature of PD, including impaired decision making and attention.^4–7^ Some patients with PD already experience this at early stages, and up to 80% develop cognitive impairment within 20 years of diagnosis as pathology progresses into cortical areas. By age 90, 80-90% are affected, and the risk of dementia is increased sixfold compared to age-matched healthy controls.^8,9^ In agreement with cognitive impairments, cortical networks become dysfunctional in PD.^10,11^ A shift in the emphasis from motor dysfunction in PD to address the understudied yet clinically significant non-motor impairments is therefore needed.

The extent of Lewy pathology in cortical neurons correlates strongly with cognitive decline and dementia in synucleinopathies other than PD.^12,13^ α-Synucleinopathy in the neocortex is prominent in Dementia with Lewy bodies (DLB) patients, where it progresses first in layer V-VI and is then detected in axon terminals.^14,15^ Cortical α-syn pathology in DLB is also correlated with intracortical dysconnectivity and cognitive impairment.^16–19^ α-Syn aggregation could therefore be directly involved in cortical impairment.

In the healthy brain, α-syn is enriched in presynaptic terminals, and the three members of this protein family shape synaptic architecture.^20^ This abundance of endogenous α-syn at synapses may make them a neuronal compartment particularly vulnerable to α-synucleinopathy. Indeed, presynaptic specializations are less abundant in persons with PD and DLB^21–23^ and increased loss of presynaptic markers is associated with higher cortical LB pathology.^24^ Furthermore, the density of dendritic spines, the postsynaptic sites of excitatory inputs, is reduced in several brain regions in PD and in models of α-synucleinopathies.^25–27^ These synaptic aberrations in synucleinopathies could contribute to cognitive impairments at later stages. Understanding the progression of these aberrations is therefore important, and studies in the striatum have shown progressive impairment of both excitatory synapses and dopamine release sites in a PD model.^28^ Yet, how Lewy pathology impacts synaptic density and morphology and the temporal and spatial patterns of synapse loss in the cortex remain to be defined.

We analysed here the molecular properties and the temporal and spatial relationships of cortical synaptopathy driven by α-syn aggregates using intrastriatal injection of α-syn pre-formed fibrils (PFFs) in mice. Synaptic α-syn aggregation and excitatory synapse loss initially occurred close to neuritic aggregates, which are predominantly axonal. We identified structural aberrations and progressive loss of VGLUT1-positive intracortical excitatory synapses, and excitatory transmission was impaired. In contrast, the less abundant VGLUT2-positive long-range inputs remained intact in the cortex. Further, the patterns of excitatory synapse loss corresponded to cortical neuroanatomical connectivity. These results, obtained through super-resolution imaging, electron microscopy, mesoscale imaging, and electrophysiology, demonstrate molecular, structural, and functional properties of synaptic pathology and support a progressive vulnerability of intracortical synapses to α-syn pathology in this model. Our findings establish a framework for understanding synaptic vulnerability to α-syn pathology and its progression in cortical circuits.

## Materials and Methods

An extended version of the Materials and Methods is available in the Supplementary material.

### Antibodies

Supplementary Table 1 provides primary antibodies and application notes for immunohistochemistry (IHC). Secondary antibodies were conjugated to Alexa dyes 488, 568, and 647 (Thermo Fisher). Isotype-specific secondary antibodies were used to detect monoclonal primary antibodies. For mesoscan imaging, the antibody against phosphorylated α-syn was directly conjugated with Alexa-647, as described in the Supplementary material, ‘Methods’ section.

### Virus

pAAV-CAG-GFP was a gift from Dr. Edward Boyden (Addgene viral prep # 37825-AAVrg; http://n2t.net/addgene:37825 ; RRID: Addgene_37825).

### Animals

3-month-old C57BL/6J mice (RRID:IMSR_JAX:000664) and 5-8 weeks old Snca^GFP^ knock-in mice (RRID:IMSR_JAX:035412)^29^ were obtained from Jackson Laboratories (Bar Harbor, ME, USA). Mice were injected with either PFF or monomer into the striatum. All animal procedures were approved by the respective University Institutional Animal Care and Use Committees and complied with the National Institutes of Health guidelines for the Care and Use of Laboratory Animals (Supplementary material, ‘Methods’ section).

### Stereotaxic injections of α-synuclein monomer and preformed fibrils

Mice were deeply anesthetized using an isoflurane vaporizer and placed on a stereotactic frame. Mice were injected with PFFs to induce α-syn pathology or α-syn monomer as a control into the right forebrain to target the dorsal striatum. PBS was injected as a control for electron microscopy. The animals were treated postoperatively with carprofen after injection and kept under observation for 72 hours after surgery (Supplementary material, ‘Methods’ section).

### Tissue processing for synapse quantification

40-50 µm thick brain slices were obtained and processed for SEQUIN imaging.^30,31^ Sections stained for SEQUIN analysis utilized a combination of antibodies targeting pre- and post-synaptic structures for the identification of synaptic loci based on trans-synaptic separation. Slides were coverslipped using the high-refractive index mounting media MWL 4-88 (Electron Microscopy Sciences) and high-precision No. 1.5H cover glass (Marienfeld) (Supplementary material, ‘Methods’ section).

### Synaptic imaging and analysis

Super-resolution images for synaptic analyses were obtained, processed, and analysed as previously described,^30,31^ using a Zeiss LSM Airyscan Microscope (Carl Zeiss Microscopy) with 63x 1.4NA objective. Analysis of layer V excitatory or inhibitory synapses was performed using Imaris (Bitplane) and MATLAB (Mathworks). Analysis of cortical mesoscans was performed using Python. Synaptic colocalization analysis of endogenous α-syn with VGLUT1 and VGLUT2 was performed using Imaris (Supplementary material, ‘Methods’ section).

### Synuclein pathology progression analysis

Perfusion-fixed brain tissues were collected from mice at 1, 3, and 6 months post-injection (MPI) of either PFF or control monomer in the striatum. Immunohistochemistry was performed on cortical sections to analyse α-syn pathology propagation over the 1-6 month time course using an antibody that detects phospho-α-syn. The extent of pathology propagation was measured as increases in phospho-α-syn intensity and inclusion size and analysed using Imaris software. Localization of neuritic phospho-α-syn to axons and dendrites was performed on tissue co-labelled with Neurofilament-H and MAP-2 and analysed using Python (Supplementary material, ‘Methods’ section).

### Electron microscopy and image analysis

Tissue samples underwent routine initial EM processing and 60 nm sections were cut using an ultramicrotome (Leica), post-stained with 2% uranyl acetate and lead citrate, and imaged with FEI Tecnai transmission electron microscope. Asymmetrical synapses were identified by a researcher blinded to experimental conditions and the area of individual synaptic vesicles was quantified using a convolution neural network trained on mouse synapses.^32^ Synaptic cleft width was measured from the edge of the pre-synaptic terminal to the centre of the post-synaptic density (Supplementary material, ‘Methods’ section).

### Electrophysiology

300 μm thick acute slices were prepared in the coronal plane from the region neighbouring to the injection site (PFF or monomer). Whole-cell recordings were carried out at RT and miniature excitatory postsynaptic currents (mEPSCs) and inhibitory postsynaptic currents (mIPSCs) were recorded for identified pyramidal cells (Supplementary material, ‘Methods’ section).

### Statistical analysis

Data were analysed using linear mixed-effects models to account for repeated measures within animals. Mesoscan data were analysed using MATLAB R2023b (fitlme). Synapse density, pre-and postsynaptic protein separation measurements, percentage colocalization, and transmission electron microscopy (TEM) data were analysed using linear mixed-effects models in SPSS (IBM Corp.). For the analysis across cortical areas, animal and brain regions were included as random effects, with treatment (control vs PFF) as a fixed effect, to avoid pseudoreplication and to ensure that individual animals served as the unit of statistical inference. TEM data showed right-skewed distributions and were preprocessed by trimming 5% extreme values and applying Box-Cox transformation before analysis using linear mixed models with mouse as a random effect and compound symmetry covariance structure to account for multiple synapses per animal. The extent of cortical synuclein pathology propagation over time was analysed using an ordinary one-way ANOVA in GraphPad Prism. Excitatory synaptic transmission in the cortex between monomer-and PFF-injected mice was compared using an unpaired t-test. Sample sizes, statistical significance set value, and data presentation details for all results are provided in the figure legends.

## Results

### α-Synuclein aggregates mark cortical excitatory synapses

The features of cortical α-synucleinopathy remain incompletely understood. We investigated them in a mouse model in which the aggregation of endogenous α-syn is templated by exogenous α-syn pre-formed fibrils (PFFs), one of the animal models of synucleinopathy.^33^ PFFs were injected into the dorsolateral striatum of wild-type mice to induce pathology in corticostriatal projection neurons (Supplementary Fig. 1A). Injection of α-syn monomer served as a control. We selected this PFF model because it replicates abundant α-syn aggregates in late-stage human PD in cortical layer V and the disruption of cortical connectivity, and allows studies of pathology progression.^26,34,35^ We analysed synaptic pathology in the frontal cortex, which is preferentially affected in PD, and focused on the secondary motor area (M2) that exhibits pathophysiological changes in synucleinopathies^1,36^ and plays roles in decision-making and cognitive control.^37,38^ α-Syn aggregates were detected immunohistochemically with antibodies against phosphorylated S129 in α-syn (p-α-syn) that detect the misfolded form.^39,40^ Unilateral striatal PFF injection resulted in the progressive α-syn aggregation in the ipsilateral cortical layer V over a six month period that was not detected after α-syn monomer control injection, as expected (Supplementary Fig. 1B-E).

We focused on layer V because of its susceptibility to human Lewy pathology.^14^ Relevant to our model, the corticostriatal projections of intratelencephalic (IT-type) and pyramidal tract (PT-type) neurons in this layer expose them to striatally injected PFFs. To investigate the early stages of cortical synaptic pathology in α-synucleinopathies, we utilized the temporal control over pathology onset provided by the striatal PFF injection model. We first analysed the frontal cortex at 1 month post-injection (MPI). We performed super-resolved SEQUIN 3D imaging of the presynaptic vesicle protein synapsin and the postsynaptic scaffold protein PSD-95, together with p-α-syn, in cortical tissue (Fig. 1A). This technique employs a multi-detector array to enhance resolution beyond the diffraction limit in all spatial dimensions. Super-resolution enables synapse identification using the criterion that these markers of pre- and post-synaptic specializations are paired in space at a distance matching ultrastructurally derived parameters, and SEQUIN imaging identified excitatory synapses as spatially paired synapsin/PSD-95 loci. p-α-Syn was frequently detected at these excitatory synaptic loci in PFF-injected mice (Fig. 1B). We measured the amount of p-α-syn within a 200 nm field around the centroid of each synaptic locus (Fig. 1C). As expected, monomer injections did not result in significant immunoreactivity of p-α-syn (Fig. 1D). In the PFF-injected cohort of mice, p-α-syn was present at synapses and extrα-synaptically (Fig. 1D; p<0.0001, linear mixed-effects model). Synaptic p-α-syn levels were increased over those observed in extrα-synaptic areas, demonstrating synaptic enrichment of p-α-syn (p<0.0001, linear mixed-effects model).

**Fig. 1.**
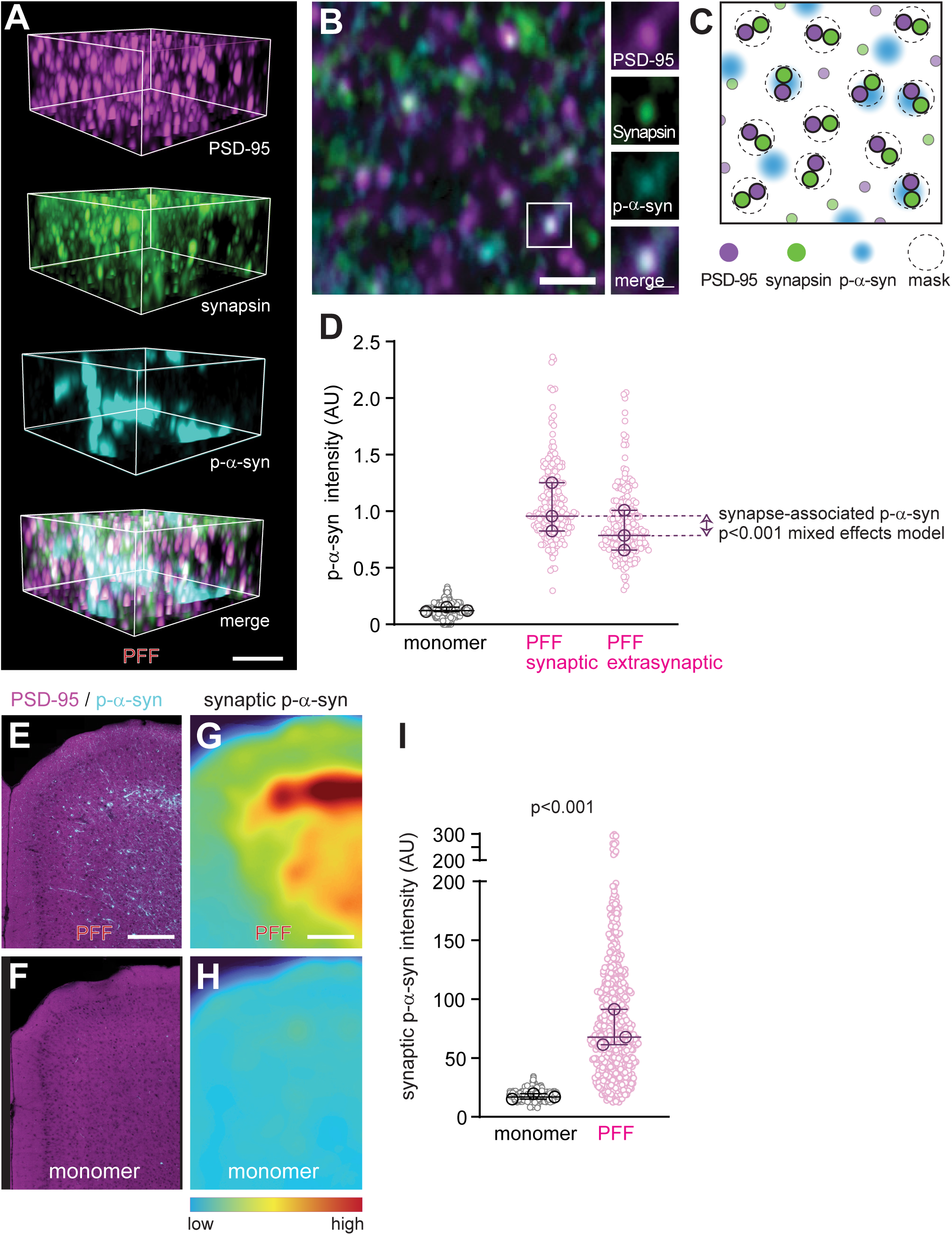
α-Synuclein aggregates in cortical excitatory synapses. **(A)** Representative 3D imaging fields immunostained for excitatory postsynaptic PSD-95 (magenta), the synaptic vesicle protein synapsin (green), and p-α-syn (cyan) in frontal cortex M2 at 1 MPI of PFFs. Scale bar, 5 µm. **(B)** High magnification 2D projection at 600 nm thickness showing the synaptic overlap of p-α-syn with spatially paired synapsin and PSD-95. Scale bar, 1 µm. **(C)** Schematic of measuring p-α-syn immunostaining in 200 nm fields centred on synaptic loci. **(D)** Quantification of cortical p-α-syn, from the area shown in (**E**), in monomer-injected control mice and synaptic and parenchymal p-α-syn in PFF-injected animals. Small circles show data from individual image tiles, and large circles show the median for each animal. Bars mark the group-wise median values of synaptic and extrasynaptic p-α-syn in PFF mice, and arrowheads depict the amount of p-α-syn that is synapse-associated. Statistical analyses were performed on image tiles using a linear mixed-effects model to account for non-independent data. (monomer, n=1019 tiles; PFF, n=196 high path area tiles; three mice per group) (**E, F**) Representative mesoscale tilescan of frontal cortex M2 after immunostaining for excitatory postsynaptic PSD-95 (magenta) and p-α-syn (cyan) at 1 MPI of PFF (**E**) or monomer (**F**). Maximum intensity projections are shown. Scale bar, 250 µm. (**G, H**) Groupwise average heatmaps of synaptic p-α-syn abundance from PFF- (**G**) or monomer-(**H**) injected animals in the area shown in (**E**) and (**F**). Scale bar, 250 µm. Synapses were first identified using SEQUIN-based measurement of pre-to-postsynaptic separation, then synaptic p-α-syn was quantified per synapse as shown in (**C**). The average local intensity of synapse-associated p-α-syn was used to generate a heatmap for each subject. Heatmaps were transformed to a common coordinate frame and averaged to produce the groupwise heatmaps in (**G**) and (**H**). **(I)** Synaptically localized p-α-syn from areas as in (**G**) and (**H**) was detected 1 MPI of PFF compared to monomer and intensity was measured. The graph presents image tile and animal data as in (**D**). Statistical analyses were performed using a linear mixed-effects model. (monomer, n=598 image tiles; PFF, n=565 tiles; three mice per group)

Mesoscans across cortical layers confirmed prominent parenchymal p-α-syn pathology in layer V of PFF- but not control-injected mice at 1 MPI (Fig. 1E and F). Super-resolved imaging of synaptic markers and p-α-syn enabled us to identify the synaptic fraction of cortical p-α-syn in the mesoscans, as visualized in heatmaps (Fig. 1G and H). This confirmed that synaptically localized p-α-syn is abundant in PFF-injected mice compared to monomer controls (Fig. 1I) (p<0.0001, linear mixed-effects model). Synapses, therefore, are cellular loci of α-syn pathology in the cortex.

### Proximity to Lewy neurite-like structures increases synaptic α-syn pathology and loss

The spatial relationship between neuritic p-α-syn aggregates and cortical synapse loss is unknown. We analysed this at 1 MPI of PFFs or α-syn monomer control to gain insights into the early stage of pathology. Synapses were identified using SEQUIN after immunostaining for synapsin and PSD-95, and aggregates immunopositive for p-α-syn, including Lewy-neurite like structures, were imaged in the same area (Fig. 2A). Measuring the intensity of p-α-syn at synapses along with the distance between synapses and the nearest Lewy-neurite like structure revealed that synaptic p-α-syn is greatest nearby such structures (Fig. 2B). To gain additional insights into the local impact of pathology on synapse loss, we quantified the density of synaptic loci within 800 nm of the perimeter of p-α-syn aggregates and analysed the extent to which proximal synaptic density was related to the relative p-α-syn immunopositivity of neighbouring neuritic aggregates. Synapse loss was most pronounced in close proximity to the neuritic structures with the highest p-α-syn labelling at this early 1 MPI stage of pathology (Fig. 2C). Structural changes were observed for the remaining synaptic loci in proximity to p-α-syn aggregates, which exhibited enlarged pre-synaptic and smaller post-synaptic puncta (Supplementary Fig. 2). These results support that proximity to neuritic α-syn aggregates increases localized intrα-synaptic α-syn accumulation and structural changes and synapse loss already at the stage of pathology onset.

**Fig. 2.**
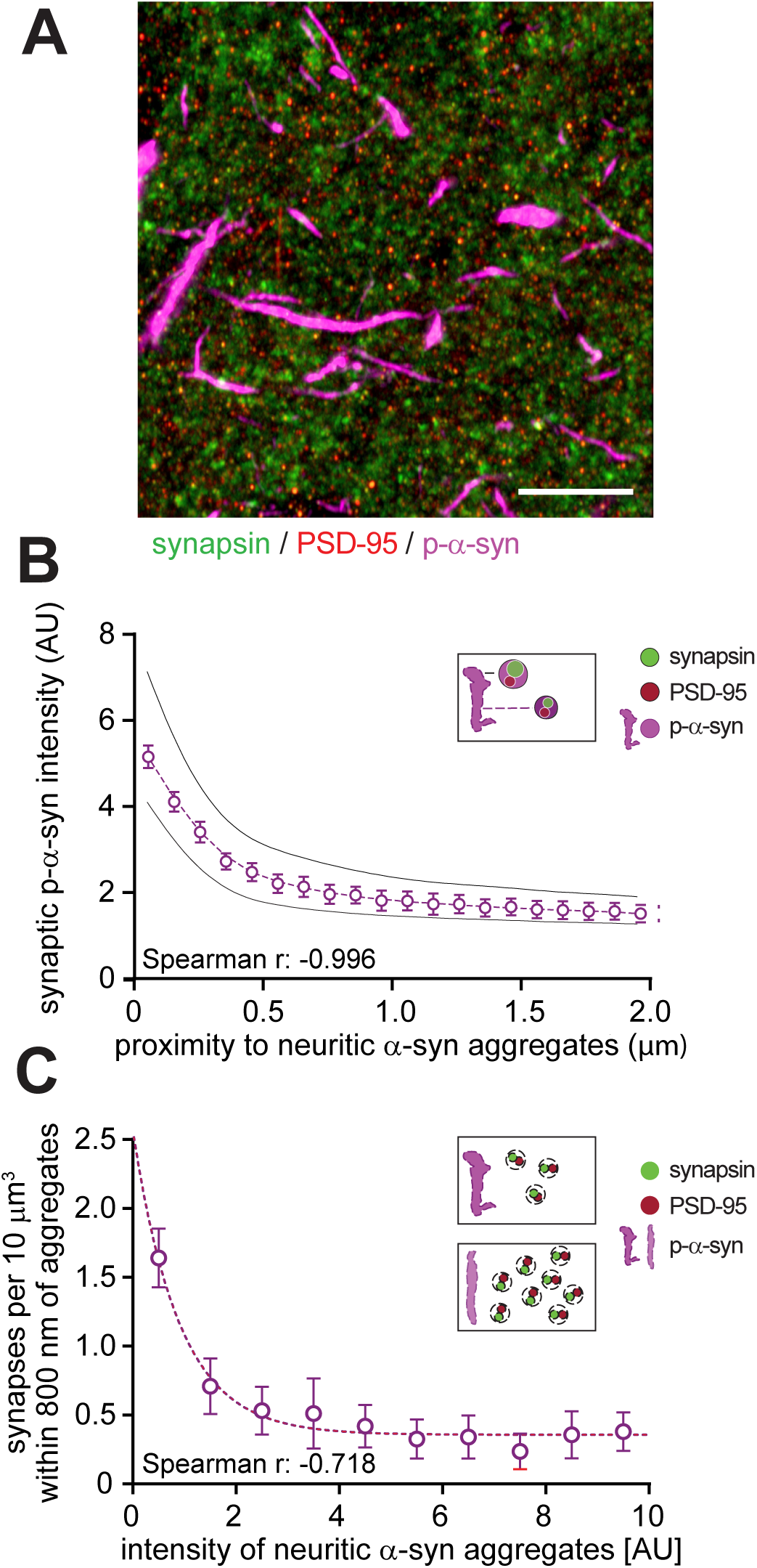
Proximity to Lewy neurite-like structures impacts synaptic pathology. **(A)** Representative image of presynaptic synapsin (green), postsynaptic PSD-95 (magenta), and p-α-syn aggregates (cyan) in the cortex 1 MPI of PFF. Scale bar, 10 µm. **(B)** Quantification of images as in (**A**) shows that synaptic p-α-syn enrichment is correlated with the distance to the nearest Lewy neurite-like p-α-syn aggregate. **(C)** Synapse density was analysed within 800 nm of neuritic p-α-syn aggregates. Aggregates were binned by the extent of their p-α-syn signal. Synapse density was lowest proximal to the aggregates with the highest p-α-syn intensity. A within-intensity-bin median calculation was used for density measurement. (**B** and **C**, monomer, n=598 image tiles; PFF, n=565; three mice per group; Spearman’s rank correlation coefficients provided in the panels)

### Lewy neurite-like structures are more frequently associated with axons than dendrites

To define the neuritic location of the cortical p-α-syn aggregates induced in our model, PFF-injected animals were co-injected with a retrograde AAV to label neurons projecting to the injection site with GFP. Tissue from these animals was obtained at 2 weeks post-injection, then immunostained for p-α-syn. Aggregates were readily detectable in GFP-positive axons in the ipsilateral corpus callosum (Fig. 3A). To quantitate p-α-syn association with axons and dendrites, ipsilateral samples from frontal cortex 1 month post PFF injection were co-labelled with the axonal marker NF-H, the dendrite marker MAP-2, and against p-α-syn (Fig. 3B and C). Analysis of super-resolution tilescans allowed imaging these tightly packed neurites (Supplementary Fig. 3A to H). This revealed that p-α-syn aggregates in the cortex ipsilateral to the injection site were more frequently associated with axons than with dendrites (Fig. 3D). Analysis of the contralateral cortex showed a similarly elevated frequency of aggregates associated with axons compared with dendrites, however, the total number of colocalized aggregates was lower (Fig. 3D). This preference for p-α-syn association with axons over dendrites disappeared when the neuritic masks were randomized and thus is not a function of axonal or dendritic label density (Supplementary Fig. 3I). As p-α-syn aggregate size increased, their relative preference for axons also increased, suggesting that molecular features of aggregates differ between neurites (Fig. 3E). These results demonstrate that neuritic p-α-syn aggregates in the PFF model preferentially associate with cortical axons, though dendrites are also affected.

**Fig. 3.**
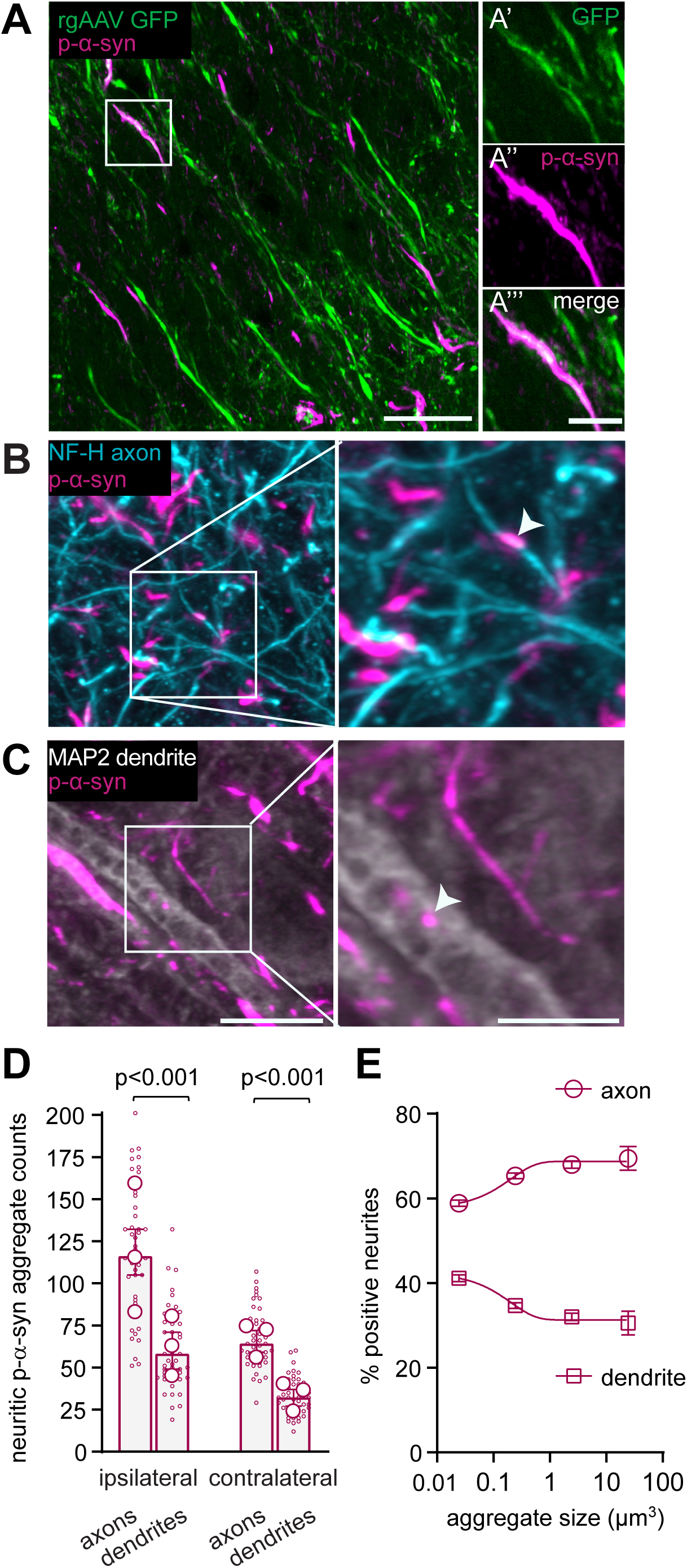
α-Synuclein aggregates are more frequently detected in axons than in dendrites. (**A**) Representative image from ipsilateral frontal cortex of a PFF injected animal showing p-α-syn aggregates and axons retrogradely labelled with GFP. Scale bar, 20 µm. **A’** Zoom of GFP labelled axon boxed in (**A**) colocalized with p-α-syn (**A’’**). **A’’’** shows merged image. Scale bar, 5 µm. (**B, C**) Representative images from ipsilateral frontal cortex 1 month after PFF injection, labelled for the axonal marker Neurofilament-H (NF-H), the dendritic marker MAP-2, and p-α-syn. Zoomed region to the right shows labelled axons from area boxed in (**B**) and p-α-syn localized to axons in this area. Zoomed region to the right shows labelled dendrites from the area boxed in (**C**) and p-α-syn localized to dendrites in this area. Scale bar in (**C**), 10 µm, also applies to (**B**). Scale bar in zoomed **(C**) field, 5 µm; also applies to zoomed (**B**). Arrowheads identify examples of p-α-syn colocalized with an axon or dendrite. **(D)** Quantification of p-α-syn localization reveals more frequent detections in axons compared to dendrites. Analysis of contralateral cortex revealed fewer total colocalizations, indicating lower overall number of p-α-syn aggregates, while a similar ratio of axon to dendrite localizations was observed. (n=16 image tiles per animal from three mice, linear mixed-effects model). **(E)** Quantification of the frequency of axonal vs dendritic localization of p-α-syn by aggregate size shows that the ratio of axonal to dendritic localization of aggregates is more biased towards axons at increasing aggregate size.

### Ultrastructural synaptic aberrations occur in the presence of cortical α-syn pathology

Given the size changes of synapses in proximity to aggregate bearing neurites at initial stages, we hypothesized that α-syn pathology may affect other structural properties of excitatory synapses that have not undergone loss. We first used SEQUIN to measure the separation of pre- and post-synaptic VGLUT1- and Homer-positive assemblies in the frontal cortex after striatal injection of PFFs vs. monomer. The Gaussian distribution of pre- and post-synaptic distances at synaptic loci showed no difference at 1 MPI (data not shown). However, at 3 MPI, when pathology is more widespread (Supplementary Fig. 1), we detected a greater separation of VGLUT1/Homer specializations after striatal injection of PFFs than after monomer (Fig. 4A-C).

**Fig. 4.**
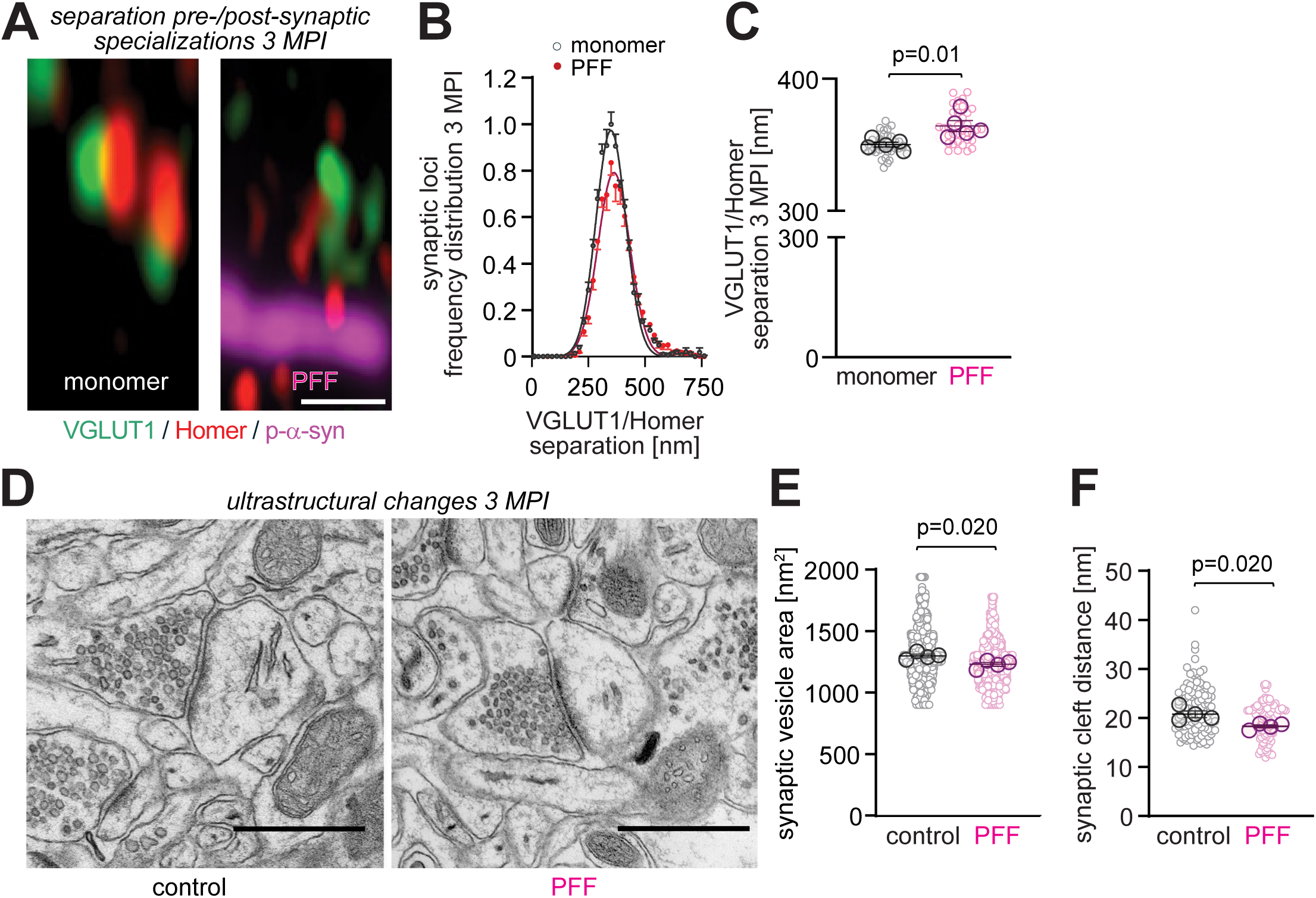
Structural synaptic aberrations in the presence of cortical α-synuclein aggregates. **(A)** Representative high-resolution Airyscan images from the frontal cortex M2 in layer V. Images were acquired 3 MPI of monomer control (left) or PFF (right) and show paired pre- and post-synaptic excitatory VGLUT1 (green) and Homer (red) specializations, with p-α-syn in magenta. Scale bar, 1 µm. **(B)** VGLUT1/Homer specializations become spatially separated in PFF- compared to monomer control-injected mice. Nearest-neighbour separation distances were determined by analysing the Gaussian distribution of synaptically paired puncta and their mean distance using SEQUIN. The distribution of synaptic loci was normalized to the amplitude of 1 for monomer-injected control mice. **(C)** The separation of spatially paired VGLUT1/Homer specializations was increased in PFF-compared to monomer control-injected mice. The graph shows separation distances for individual synaptic pairs after nearest-neighbour analysis. (**B** and **C**, n=five animals/group, six-ten slices per animal) **(D)** Ultrastructural analysis of synaptic morphology in mice injected with control vs. PFF at 3 MPI in M2. Representative transmission electron microscopy (EM) images of synapses in the frontal cortex are shown. Scale bar, 600 nm. **(E, F)** Excitatory synapses undergo ultrastructural changes. Analysis of EM images as in (**D**) determined that synaptic vesicle area (**E**) and synaptic cleft width **(F**) were reduced. (E, control, n=8810 synaptic vesicles; PFF, n=7576; four mice per group; F, control, n=345 synapses; PFF, n=283; four mice per group). Results are presented as mean ± SEM, where SEM is calculated from mouse means. Small circles represent individual image measurements, whereas large circles indicate the mean per mouse. Group differences in (**C**) were assessed using a linear mixed-effects model, with group as a fixed effect and mouse as the unit of inference. Data in (**E**) and (**F**) were analysed after Box-Cox transformation using linear mixed models with synapses nested within each mouse. Statistical significance was defined as p<0.05.

To assess the effects of α-syn pathology on synaptic ultrastructure in the cortex, we employed transmission electron microscopy (TEM) (Fig. 4D). Mice were perfused for TEM at 3 MPI. Aberrations were found in multiple synaptic compartments in PFF vs. control-injected mice. Presynaptically, the area of individual synaptic vesicles (SV) was moderately but significantly reduced in PFF-injected mice compared to controls (Fig. 4E). Analysis of inter-SV distance showed a minor reduction in PFF-injected vs. control mice that reached significance, while the number of docked SV and PSD length were unchanged (Supplementary Fig. 4). Further, the synaptic cleft distance was shortened (Fig. 4F). This indicated parallel structural disruptions, impacting trans-synaptic cleft geometry on a scale of 20 nm, as measured by TEM, and a long-range redistribution of pre- and post-synaptic protein assemblies on a scale of 300-400 nm, as measured by SEQUIN imaging (Fig. 4B and C). These results demonstrate broad structural changes in cortical synapses in the presence of α-syn pathology.

### Intracortical excitatory synapses are progressively lost in the presence of α-synuclein pathology while other cortical synapse types are spared in the striatal seeding model

Misfolded α-syn deposition gradually increases across the brain in PD, including in the neocortex.^41,42^ Our PFF model replicated the increase of cortical α-syn aggregates in maturity and size over 6 months (Supplementary Fig. 1).^40,43^ To analyse progressive synaptic pathology, excitatory synapses were detected by co-immunostaining for presynaptic VGLUT1, a marker of intracortical synapses, and the excitatory postsynaptic scaffold protein Homer, followed by SEQUIN analysis of spatially paired VGLUT1/Homer loci. We investigated whether the intrastriatal injection of PFFs and the subsequent formation of α-syn aggregates in layer V affected synapse density across stages of pathology. No statistically significant difference in the overall density of intracortical VGLUT1-positive excitatory synapses was detected at 1 MPI of PFFs, the first time point analysed, compared to monomer-injected control mice (Fig. 5A and B). This agreed with the highly localized synapse loss found at 1 MPI only in the immediate proximity of neuritic aggregates (Fig. 2C). At 3 MPI, however, a significant loss of VGLUT1-positive excitatory synapses was measured in the frontal cortex in PFF-injected compared to monomer controls (Fig. 5C and D). This reduction in the density of VGLUT1/Homer-positive synapses persisted at 6 MPI (Fig. 5E and F). Unlike at 3 MPI, the distances of pre- and post-synaptic assemblies were unchanged at 6 MPI of PFFs (Supplementary Fig. 5). This indicated that synapse loss at 3 MPI involved structural destabilization, which was no longer observed by 6 MPI. Together, α-syn pathology in the cortex after striatal PFF injection is associated with a progressive loss of intracortical VGLUT1-positive excitatory synapses. To further assess excitatory synapse vulnerability, we performed SEQUIN imaging of VGLUT2-positive long-range inputs to layer V at 3 MPI. These predominantly thalamocortical projections^44,45^ showed no changes in density (Fig. 5G and H) or structural properties (Supplementary Fig. 6) in PFF- vs. monomer-injected mice. Molecular factors related to presynaptic identity hence can contribute to synapse type-specific vulnerabilities, while it also needs to be considered that the striatal seeding paradigm exposes select cortical neuron types to pathology.

**Fig. 5.**
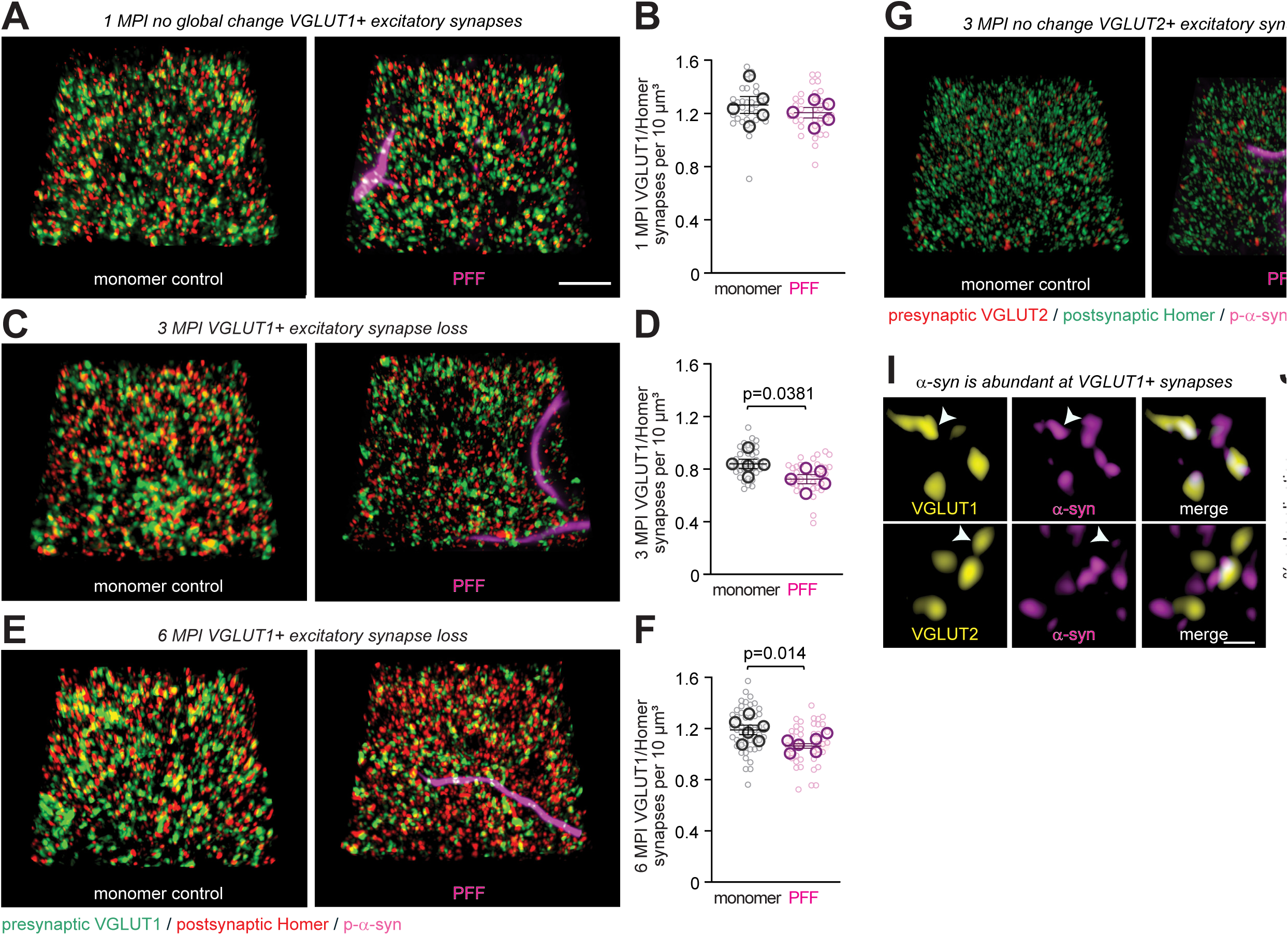
Intracortical VGLUT1-positive excitatory synapses but not long-range VGLUT2-positive inputs are progressively lost after α-synuclein pathology appears and are distinguished by α-synuclein content. **(A)** Representative 3D high-resolution immunostainings acquired by Airyscan imaging in layer V of frontal cortex, secondary motor area M2. Tissue was immunostained for excitatory presynaptic VGLUT1 (green) and excitatory postsynaptic Homer (red). VGLUT1 marks intracortical synapses. Staining for phospho-α-syn (p-α-syn, magenta) detects α-syn aggregates. Images were acquired 1 month post-injection (MPI) of monomer (left) or PFF (right) and processed for SEQUIN analysis. Scale bar, 5 µm. **(B)** Density of synapses marked by spatially paired VGLUT1 and Homer specializations was unchanged at 1 MPI of monomer vs. PFF. The density from the frequency distribution of nearest-neighbour VGLUT1/Homer pairs was calculated from images as in (**A**). (n=five animals/group, six-eight slices per animal) **(C)** Airyscan imaging of VGLUT1 and Homer with staining for p-α-syn was performed in the frontal cortex layer V at 3 MPI of monomer or PFF. **(D)** Excitatory synapse loss occurred at 3 MPI of PFF compared to monomer. Quantification as in (**B**). (n=five animals/group, six-ten slices per animal) **(E)** Airyscan imaging of VGLUT1 and Homer with staining for p-α-syn was performed in the frontal cortex layer V at 6 MPI of monomer or PFF. **(F)** Excitatory synapse loss persisted at 6 MPI of PFF compared to monomer. Image quantification as in (**B**). (n=six animals/group, six-eight slices per animal) **(G)** Representative 3D high-resolution images obtained by Airyscan microscopy after immunostaining for excitatory presynaptic VGLUT2 (red) and postsynaptic Homer (green) in frontal cortex M2. VGLUT2 marks long-range synaptic inputs. Staining for p-α-syn detected α-syn aggregates (magenta). Images were obtained 3 MPI of monomer control (left) or PFF injection (right). Scale bar, 5 µm. **(H)** Cortical α-syn aggregates do not impact the density of VGLUT2/Homer synapses. Synapse density was calculated after SEQUIN analysis of images as in (**G**) from the frequency distribution of nearest-neighbour VGLUT2 and Homer pairs. (n=five animals/group, six-eight slices per animal) **(I)** Representative high-resolution immunostaining images showing presynaptic VGLUT1 (top row), VGLUT2 (bottom), and endogenous α-syn in the cortex of control mice at 3 MPI. Arrowheads mark examples of high α-syn at VGLUT1 and low amounts at VGLUT2 sites. Scale bar, 0.5 µm. **(J)** α-Syn colocalized robustly with VGLUT1-positive excitatory sites quantified from images as in (**I**), but colocalization with VGLUT2-positive terminals was minimal. (n=four each group, six-eight slices per animal) Results are presented as mean ± SEM, calculated from mouse means. Small circles represent individual image measurements, whereas large circles indicate the mean per mouse. Group differences were assessed using a linear mixed-effects model, with group as a fixed effect and mouse as the unit of inference. Statistical significance was defined as p<0.05.

To identify molecular properties underlying differential vulnerability, we measured total α-syn levels in control mice. This determined its enrichment at VGLUT1-positive cortical terminals, while VGLUT2-positive terminals contained very low α-syn amounts (Fig. 5I and J), consistent with a prior report.^46^ These results demonstrate that long-range VGLUT2-positive inputs are spared to cortical α-synucleinopathy in this model, in contrast to the vulnerability of intracortical VGLUT1-positive synapses, and suggest that differences in endogenous α-syn levels predispose synapse subtypes to loss.

We extended our analysis to inhibitory synapses to assess the impact of α-syn aggregates on the cortex, using immunostaining for presynaptic VGAT and postsynaptic Gephyrin. SEQUIN imaging showed no effect of PFF injection on inhibitory synapse density compared to monomer at 3 or 6 MPI (Supplementary Fig. 7). The preservation of inhibitory synapses agrees with a role of endogenous synaptic α-syn in vulnerability, as it is consistent with the lower α-syn expression and lower extent of Lewy inclusion burden in cortical inhibitory neurons.^1,47^

### α-Synuclein pathology impairs excitatory transmission in the cortex

The aberrations in excitatory synapse number and structure we observed in the presence of α-syn pathology may impact synaptic transmission. We tested this by preparing acute cortical slices 5-6 weeks after striatal injection of PFF vs. monomer control and performed whole-cell voltage clamp recordings from pyramidal cells located within the region of expected pathology (Fig. 6A). Analysis of mEPSCs showed that there was no significant difference in the mean amplitude (Fig. 6B). However, the mean inter-event interval of mEPSCs was significantly higher for PFF-injected mice compared to monomer-injected controls (Fig. 6C). Analyses of mIPSCs showed no significant differences between groups for either measure (Fig. 6B and C). Excitatory neurotransmission in the cortex is hence impaired under conditions of α-syn pathology.

**Fig. 6.**
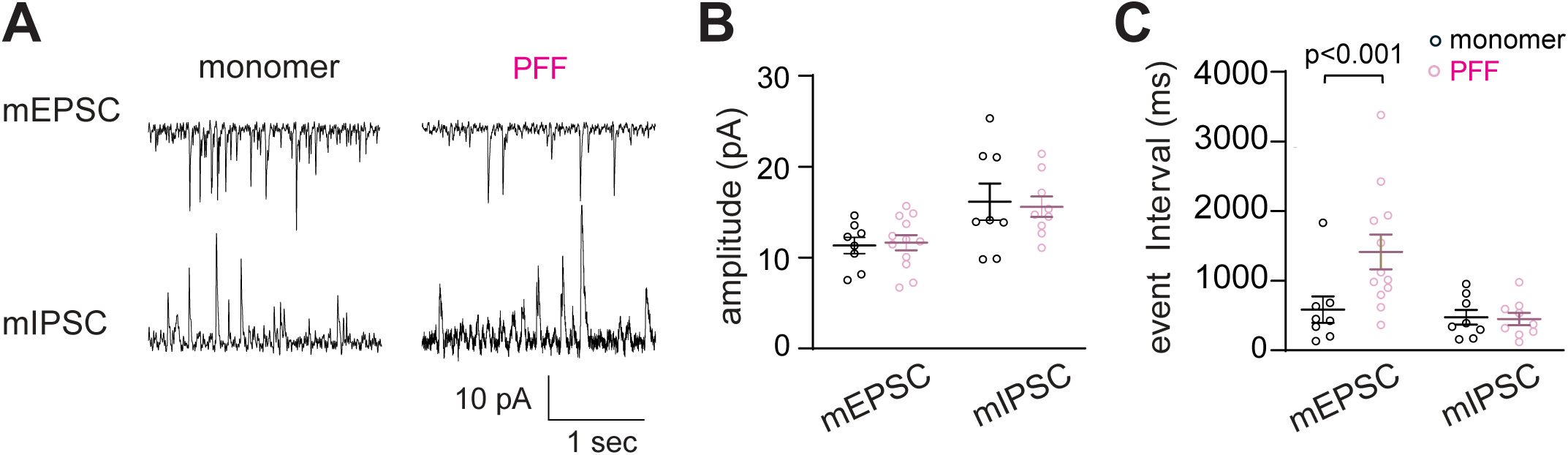
Reduction in frequency of miniature excitatory synaptic currents in mice with PFF injection. **(A)** Traces showing miniature excitatory synaptic currents (mEPSCs, top) and miniature inhibitory currents (mIPSCs, bottom) recorded from pyramidal cells in monomer (left) or PFF (right) injected mice in layer II/III at 5-6 weeks post-striatal injection. **(B)** Scatter plots of current amplitudes for mEPSCs and mIPSCs. No difference was observed in the event amplitude of mEPSCs or mIPSC between monomer- and PFF-injected mice. **(C)** Scatter plots of inter-event interval for mEPSCs and mIPSCs from monomer and PFF-injected mice. Inter-event interval was significantly higher for mEPSCs from PFF-injected mice (n=12 cells from six mice) than after monomer injection (n=eight cells from three mice). No difference was seen in the inter-event interval for mIPSCs. (monomer, n=eight cells from three mice; PFF, n=nine cells from six mice) Results are presented as mean ± SEM and considered statistically significant with p<0.05, after an unpaired t-test.

### Synaptic pathology and excitatory synapse loss are correlated across connected cortical layers

We next investigated whether intracortical aberrations also occurred in other cortical layers in the presence of p-α-syn pathology in layer V. Specifically, we focused on the relationship with layer II/III, as these neurons primarily project to layer V.^48^ Pathology was readily detectable in layers V and II/III at 1 MPI of PFFs (Fig. 7A). Using cortical mesoscans, synaptic loci were identified with SEQUIN in layers V and II/III of the frontal cortex ipsilateral to the striatal injection site. PFF-injected animals exhibited a significant loss of synaptic loci in both layers V and II/III compared to monomer-injected control mice (Fig. 7B). Analysis of individual image tiles from layer V or layers II/III showed a loss of synapses in either layer in PFF-injected mice, an effect that was more pronounced in layer V (Fig. 7C). Although a contribution of projections from layer II/III neurons to the striatum cannot be ruled out, these findings agree with a propagation of synaptic pathology across interconnected cortical layers.

**Fig. 7.**
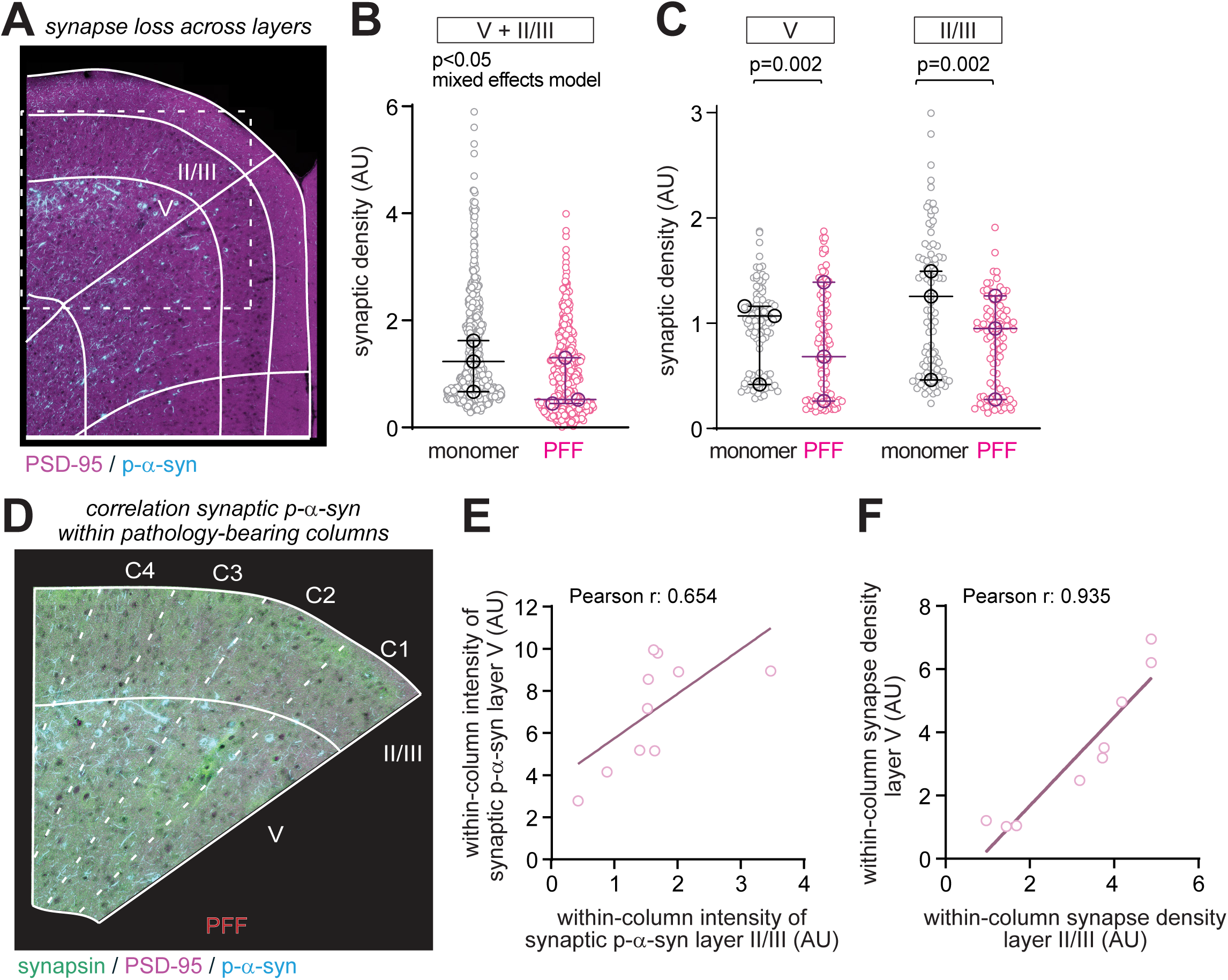
Synaptic pathology and excitatory synapse loss are correlated across cortical layers. **(A)** Representative mesoscan from a PFF-injected animal showing PSD-95 (magenta) and p-α-syn (cyan). Section-specific registration to the Allen Brain Atlas is shown in solid lines. Images were acquired in secondary motor cortex area M2 and anterior cingulate. (**B)** Mixed effects model analysis of synapse density data of combined layers V and II/III from cortical mesoscan images at 1 MPI as in (**A**). (layer V: monomer, n=304 tiles, PFF, n=285 tiles; layer II/III: monomer, n=191 tiles, PFF, n=185 tiles; from three animals per group) (**C)** Quantification of individual synaptic densities in cortex area M2 layer V and layer II/III of monomer- and PFF-injected animals from images as in (**A**). (layer V: monomer, n=112 tiles; PFF, n=110 tiles; layer II/III: monomer, n=100 tiles; PFF, n=93 tiles; three animals per group; Mann-Whitney test on individual layers) (**D**) Representative image from a PFF-treated animal at 1 MPI showing presynaptic synapsin (green), PSD-95 (magenta) and p-α-syn (cyan). Section-specific registration to the Allen Brain Atlas is shown in solid lines. The region shown corresponds to the boxed region in (**A**). (**E**) Quantification of synaptic p-α-syn immunoreactivity in layer II/III and layer V of PFF-injected animals shows that it is correlated within individual cortical columns. Images as in (**D**) were analysed. (**F**) Strong correlation of synaptic density in layers II/III and V within cortical columns after PFF injection. Images as in (**D**) were analysed. (**E** and **F**; n=3-4 cortical columns per animal from 3 animals; Pearson correlation coefficients provided in the panels)

To further assess the synaptic pathology within cortical circuits, we investigated whether the extent of pathology and synaptic density were correlated within cortical columns. We observed a strong positive correlation between the distribution of synaptic p-α-syn immunoreactivity in layers V and II/III of individual columns at 1 MPI of PFFs (Pearson r: 0.654, p=0.0403) (Fig. 7D and E). Notably, this relationship did not extend to total p-α-syn in layers V and II/III of individual columns (Pearson r: 0.469, p=0.1720; data not shown), suggesting that synaptic localization is a better predictor of p-α-syn pathology within cortical columns. A strong correlation was observed for synapse density in layers V and II/III within the same column (Pearson r: 0.935, p<0.0001) (Fig. 7F). The extent of both synaptic p-α-syn pathology and synapse density in presence of pathology is therefore proportional between interconnected layers V and II/III in the same column. Synaptic density in layer 5 and layer 2/3 in the presence of pathology remained correlated at 6 MPI (Supplementary Fig. 8). These results support a role of connectivity patterns between cortical layers in the propagation of synaptopathy induced by α-syn aggregates.

### Contralateral pathology and cortical synapse loss support a vulnerability of IT neurons

Finally, we aimed to characterize the vulnerability of neuron subtypes in layer V based on their projection properties. In this layer, IT-type neurons project bilaterally to the striatum, unlike ipsilaterally projecting PT-type neurons^48^ (Fig. 8A). Using the unilateral injection approach, we first determined the extent of α-syn pathology in the cortical hemisphere contralateral to the striatal injection site, which would only involve IT-type neurons. We observed a substantial presence of p-α-syn immunopositive aggregates in the contralateral layer V at 6 MPI after unilateral injection of PFFs into the dorsal striatum (Fig. 8B), replicating earlier data.^40^ We next tested whether this contralateral pathology is associated with changes in the number of excitatory synapses. SEQUIN imaging of VGLUT1/Homer-positive synapses in layer V contralateral to the striatal injection site showed a significant loss of their synaptic loci at 6 MPI (Fig. 8C and D). These results align with the vulnerability of IT neurons, which project bilaterally, and agree with the dysregulated gene expression in IT neurons bearing PD pathology.^1^ This supports the notion that an imbalance of corticostriatal systems, involving IT versus PT-type neurons, contributes to neurodegenerative and other brain disorders,^48^ and that synaptic disruption is integral to this imbalance.

**Fig. 8.**
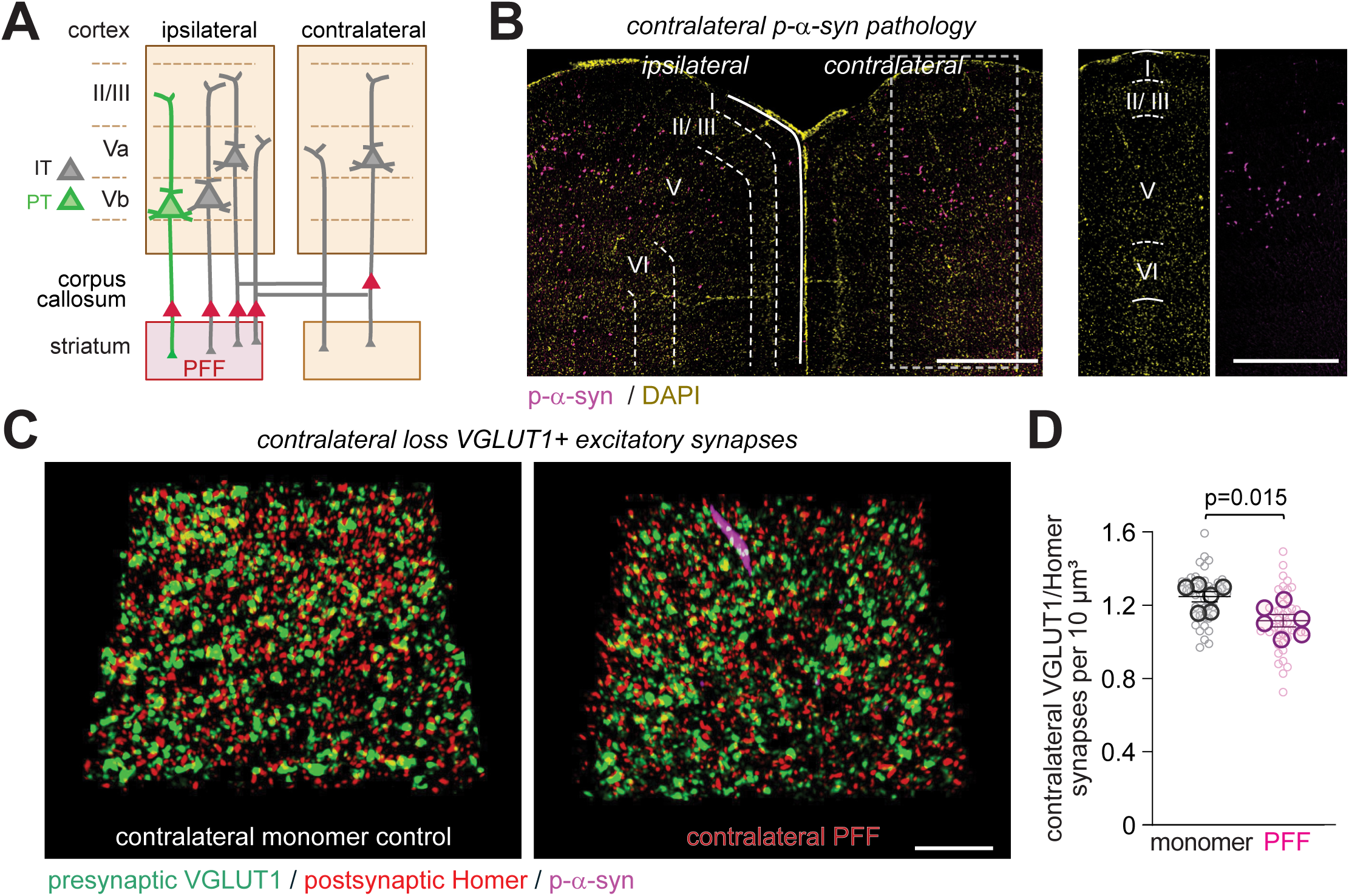
Contralateral pathology and synapse loss support vulnerability of IT neurons in layer V. (**A**) Schematic of IT and PT neuron connectivity. IT neurons (grey) project ipsilaterally to the striatum and to the contralateral striatum and cortex. PT neurons (green) project only ipsilaterally to the striatum. Red arrows indicate the direction of α-syn pathology spread. Adapted from Shepherd (2013). ^48^ (**B**) Left, representative tiled overview image of p-α-syn immunostaining (magenta) in layer V of cortical hemispheres ipsilateral and contralateral to the striatal PFF injection site at 6 MPI, co-stained with DAPI (yellow). Right, p-α-syn–positive aggregates in contralateral layer V at 6 MPI, corresponding to the dotted square in the contralateral cortex shown in the overview. Scale bars, 500 µm. (**C**) Representative 3D high-resolution images from layer V of the frontal cortex M2 contralateral to the striatum unilaterally injected with PFFs. Images were acquired using Airyscan. Tissue was immunostained for excitatory presynaptic VGLUT1 (green), postsynaptic Homer (red), and p-α-syn (magenta). Images were acquired at 6 MPI following injection of monomer (left) or PFF (right) and processed for SEQUIN analysis. Scale bar: 5 µm. (**D**) Excitatory synaptic density, defined by spatially paired VGLUT1 and Homer specializations, was reduced in the contralateral frontal cortex after ipsilateral PFF injection compared to monomer. Quantification was performed from images as in (**C**) by analysing the frequency distribution of nearest-neighbour VGLUT1/Homer pairs. (n=six animals/group, six–eight slices per animal) Results are presented as mean ± SEM, calculated from mouse means. Small circles represent individual image measurements, whereas large circles indicate the mean per mouse. Group differences were assessed using a linear mixed-effects model, with group as a fixed effect and mouse as the unit of inference. Statistical significance was defined as p<0.05

## Discussion

This study integrates analyses across scales to reveal four insights into synaptic vulnerability in α-synucleinopathies. First, α-syn aggregation at cortical synapses is a hallmark of synucleinopathies and more strongly correlated with their loss than overall α-syn pathology. Second, cortical Lewy neurite-like pathology is predominantly associated with axons and drives early α-syn aggregation at adjacent synaptic sites, identifying a mechanism that initiates cellular pathology. Third, intracortical excitatory synapses exhibit progressive loss, whereas other synaptic populations are spared. This selective vulnerability tracks endogenous synaptic α-syn content. Fourth, the architecture of cortical circuits, together with neuron type-specific vulnerabilities, aligns with the progression of synaptic pathology across layers and hemispheres.

Our results identify synaptic α-syn aggregation as a hallmark of cortical α-synucleinopathy, indicating that synapses are early subcellular targets. Excitatory synapses near Lewy neurite-like aggregates showed p-α-syn immunopositivity already at 1 MPI, with proximal neuropil volumes exhibiting the earliest loss of synapses. Further, presynaptic terminals near aggregate-bearing neurites are enlarged. It remains unclear whether this synaptic pathology and loss proximal to neuritic α-syn aggregates result from the generation of a toxic microenvironment, which can be tested in future studies. Regarding the neuritic localization of the aggregates, the p-α-syn-positive neurites analysed here in the cortex are unlikely to arise from striatal inputs, as medium spiny neurons project predominantly to the globus pallidus and substantia nigra without known direct neocortical projections.^49–51^ These neurites therefore represent projections of corticostriatal neurons, in which pathological α-syn has propagated retrogradely from the striatal injection site. This agrees with the predominantly axonal distribution we report for neuritic α-syn aggregates.

Notably, we identified temporal patterns of vulnerability of cortical excitatory synapses, in which early structural disruption may lead to loss. By 3 MPI, widespread loss of VGLUT1-positive synapses in layer V emerged, accompanied by a separation of pre- and post-synaptic assemblies away from synaptic contact sites, indicating structural disruption. While VGLUT1-positive synapse loss persisted at 6 MPI, synaptic separation normalized. These temporal progression data may be valuable for identifying windows for disease-modifying interventions. Synapse loss affected VGLUT1-positive intracortical synapses but spared VGLUT2-positive long-range synapses despite sharing postsynaptic Homer-positive specializations. These data point to the presynaptic compartment as a determinant of vulnerability. Our results are consistent with prior findings demonstrating that VGLUT2-positive synapses in the striatum and basolateral amygdala remain unaffected in the presence of α-syn pathology.^46,52^

At least part of the synapse type-specific risk could be due to differences in the presynaptic expression of endogenous α-syn, which we found to be abundantly expressed at vulnerable VGLUT1-positive cortical terminals but barely detectable at VGLUT2-positive thalamic inputs in the cortex, as reported.^46^ In additional support of presynaptic susceptibility, we found that synaptic vesicles at excitatory cortical synapses were smaller when α-syn pathology was present, which may reflect an impairment in the endogenous function of α-syn in regulating SV endocytosis and recycling.^53–55^ These findings suggest that presynaptic α-syn content contributes to synapse-type-specific vulnerability through templated pathology progression. A limitation of the striatal seeding approach is that pathology propagates through defined anatomical connections, constraining exposure to neurons with striatal connectivity and their targets within intracortical circuits. This impacts the ability to fully resolve synapse type-specific vulnerability across the broader cortex.

The synaptic changes we describe are in line with human post-mortem evidence, though direct comparisons need to consider differences in disease stage and synapse-type-specific marker. In DLB cortex, p-α-syn aggregates localize at enlarged synaptophysin-positive presynaptic terminals across all cortical layers, affecting both pre- and post-synaptic compartments.^15^ In PD, cortical presynaptic integrity is reduced as measured by SV2A-positive site density,^22^ and PD, PDD and DLB cases show decreased synaptophysin and SV2A densities correlating with Lewy body burden.^24^ Further, work on dopaminergic release sites in early PD shows that p-α-syn is enriched at these sites and linked to axonal terminal loss.^56^ In our model, p-α-syn localizes to excitatory synapses, which is associated with reduced synaptic density and enlarged presynaptic puncta near Lewy neurites, consistent with these human observations.

The spatial patterns of vulnerability to cortical α-synucleinopathy we report agree with the model of α-syn pathology propagation between synaptically connected neuronal populations.^57–59^ We utilized the temporal and spatial control afforded by the PFF model to gain insights into how cortical circuits dynamically pattern synaptic pathology. Striatal PFF injection allowed not only the tracking of pathology progression to cortical layer V but also the detection of α-syn aggregates in layer II/III, consistent with earlier reports.^1,34^ This was accompanied by excitatory synapse loss in both layers, and interconnected layers V and II/III within the same columns exhibited a correlated loss of synapses. This suggests spread within columns as a pathological mechanism, though this is not directly tested here, and a possible contribution of projections from layer II/II to the striatum needs to be considered. Additional insight into spatial progression patterns is provided by the loss of synapses in the contralateral hemisphere at the latest 6 MPI stage analysed here. Moreover, synapse loss within intra-columnar circuits was preferentially associated with synaptic p-α-syn, as we observed in the vicinity of neuritic aggregates. This establishes that synaptic, rather than non-synaptic, α-syn aggregates represent a principal site of synaptopathy.

Layer V intratelencephalic neurons are selectively vulnerable to α-syn pathology in both human post-mortem tissue and the striatal PFF model,^1^ and the synaptic loss reported here may therefore reflect a pattern relevant to human disease. Our analysis of synapse loss in layer V, contralateral to the striatal PFF injection site, supports this notion that disease progression follows IT neuron connectivity, which project bilaterally to the striatum. This is consistent with a recent study showing that α-syn pathology is correlated with a loss of dendritic spines in IT neurons in the secondary motor cortex.^34^ Vulnerable IT neurons are integral to neural circuits spanning intracortical layers and contralateral projections, thereby establishing a circuit framework for the progression of pathology. Our experimental design does not allow a direct comparison of IT neurons with other cortical neuron types, such as pyramidal tract (PT) neurons that project beyond the telencephalon. Subsequent studies can map synapse vulnerability onto different cortical neuron types to determine whether α-syn pathology selectively targets IT neurons or affects broader populations, with consequences for cortical circuit integrity and risk in Parkinson’s disease.

The impairments in synapse number and structure in the presence of cortical α-synucleinopathy may contribute to functional deficits, as observed in other regions.^60,61^ We identified previously unreported nanoscale synaptic features, including synaptic cleft dimensions and inter-vesicle distances, together with an enlargement of presynaptic sites reminiscent of findings in the amygdala in presence of pathology.^60^ Functionally, we observed physiological deficits in the cortex of our model, characterized by a reduction in miniature EPSC frequency. This may not only reflect the loss of intracortical synapses but also pathological aggregate interference with α-synuclein’s endogenous roles in synaptic vesicle endocytosis. Our findings align with *in vitro* studies showing that PFFs alter excitatory synaptic activity in a time- and dose-dependent manner^62^ and that α-syn aggregation affects excitatory synaptic transmission.^27,63,64^ This work also complements recent PFF seeding studies analysing cortical and corticostriatal synaptic vulnerability. Pérez-Acuña and colleagues^27^ reported a loss of VGLUT1-positive cortico-cortical synapses at layer 5 and reduced spontaneous EPSC frequency without amplitude change at 20 weeks post-PFF injection, and Brzozowski et al.^52^ showed reduced VGLUT1-positive corticostriatal synapses with impaired evoked EPSCs at six weeks post-striatal PFF injection. Our recordings align with these excitatory deficits. The preserved inhibitory transmission in our study, evidenced by unchanged miniature IPSC frequency and amplitude and stable inhibitory synapse density at three and six months post-injection, contrasts with the reduction in GAD67-positive neurons in layer 5/6 of the somatosensory cortex at nine months post-injection reported by Blumenstock et al.,^65^ when α-syn pathology had spread across all cortical layers, which suggests inhibitory circuit disruption as a later disease feature. The circuit hyperactivity reported in this study ^65^ may in part be due to an emerging intrinsic hyperexcitability of IT neurons.^34^ We did not test this late stage because we focused on progression during earlier stages of pathology.

Together, our results reveal the disruption of intracortical excitatory connectivity and its association with synaptic α-syn pathology as hallmarks of α-synucleinopathy, demonstrating that synapse loss follows molecular and circuit-based rules. The synapse changes we report here parallel the decreased functional connectivity in cognitively impaired PD patients^66,67^ and altered cortical circuit connectivity in DLB. ^18^ Our findings provide insights into neuronal and subcellular basis of synaptopathy, which is incompletely understood in α-synucleinopathies.^33^ This highlights synaptopathy as central to progressive cortical network decline in PD^68,69^ and provides a synaptic framework for understanding how α-syn pathology translates into circuit-level dysfunction.

## Conclusion

This study defines cortical synaptopathy as a hallmark of α-synucleinopathies, revealing previously unrecognized molecular, temporal, and spatial determinants of synaptic vulnerability. Our analyses across scales identify intracortical excitatory synapses as early sites of α-syn aggregation in the prefrontal cortex. Moreover, this work provides insights into molecular factors of excitatory synapse vulnerability, with high synaptic α-syn abundance, as in intracortical synapses, as a risk factor. These results support the notion that synaptic α-syn aggregation initiates synapse disorganization and loss. Our physiological data, obtained at an early stage, demonstrate functional consequences of synaptic p-α-syn accumulation.

The progression from early synaptic α-syn aggregation driven by local Lewy neurites to the later widespread loss of excitatory synapses within cortical layers defines discrete pathological stages on a circuit level. A conclusion from our circuit analysis is that cortical synaptopathy targets IT neurons and their interconnected cortical layers, as well as their contralateral projections. Together, the molecular determinants of synaptic vulnerability, combined with circuit-based progression patterns, define structural and functional impairments through which α-synucleinopathies affect synapses and can progressively disrupt cortical connectivity.

## Data Availability

All resources used and generated in this study, including datasets, image sets, protocols, reagents, organism strains, and software/code, have been compiled in the Key Resources Table (KRT) and deposited on Zenodo (DOI: 10.5281/zenodo.19721669).

## Acknowledgments

We want to acknowledge Dr. Tina Matos for her contribution as our ASAP Project Manager, Dr. Sreeganga Chandra and Dr. Michael Henderson (Van Andel Institute) for input on project design and the manuscript, and Pranav Shedge for technical support. We thank Dr. Xinran Liu, Department of Cell Biology, Yale School of Medicine, for enabling the EM work as Director of the Electron Microscopy Core Facility, and Dr. Richard Kennedy (University of Alabama at Birmingham) for expert advice on statistical analyses.

## Author Contributions

All authors conceived approaches. S.S. performed mouse injections and performed and analysed immunohistochemical studies; A.D.S. performed immunohistochemical studies, optimized imaging approaches, and developed and optimized image quantification; D.P.A. performed retrograde AAV and PFF co-injections to label cortical neurons projecting to the injection sites and conducted immunohistochemical studies; J.G. and L.M., electrophysiological recordings; J.E.R. and D.L.R., EM quantifications; J.E.R., D.L.R., and V.S., preparation of PFFs; L.A.V.-D., supervised V.S., J.E.R., D.L.R., and provided manuscript input; M.J.H., supervised J.G. and L.M. and provided manuscript input; T.T.K., supervised A.D.S. and provided manuscript input; T.B., supervised S.S. and D.P.A. and wrote the manuscript.

## Supplementary Material

A detailed Methods and Materials section, along with a table and figures, is available as Supplementary material.

