## Supplementary Materials for "Progressive Vulnerability of Cortical Synapses in α-Synucleinopathy"

### Methods

#### Antibodies

| Target | Vendor | Cat. No. | RRID | Host | Clonality | Application IHC |
|---|---|---|---|---|---|---|
| VGLUT1 | Synaptic Systems | 135 304 | AB_887878 | Guinea Pig | Polyclonal | 1:200<br>(48 h, RT) |
| VGLUT2 | Millipore | AB2251 | AB_2665454 | Guinea Pig | Polyclonal | 1:300<br>(48 h, RT) |
| Synapsin 1/2 | Synaptic Systems | 106004 | AB_1106784 | Guinea Pig | Polyclonal | 1:1000<br>(72 h, RT) |
| Homer | Synaptic Systems | 160 003 | AB_887730 | Rabbit | Polyclonal | 1:250<br>(48 h, RT) |
| PSD-95 | Invitrogen | 51-6900 | AB_2533914 | Rabbit | Polyclonal | 1:200<br>(72 h, RT) |
| VGAT | Synaptic Systems | 131 003 | AB_887869 | Rabbit | Polyclonal | 1:250<br>(36 h, 4 °C) |
| Gephyrin | Synaptic Systems | 147 111 | AB_887719 | Mouse | Monoclonal IgG1 | 1:250<br>(36 h, 4 °C) |
| pS129 α-syn | BioLegend | 825701 | AB_2564891 | Mouse | Monoclonal IgG2a, κ | SEQUIN, 1:2200<br>(48 h, RT for co-stain VGLUT/Homer; 36 h, 4 °C co-stain VGAT/Gephyrin)<br>Mesoscans, 1:500 (72 h, RT) |
| α-syn | BD Biosciences | 610787 | AB_398108 | Mouse | Monoclonal IgG1 | 1:1000 (48 h, RT) |
| NF-H | Millipore | AB5539 | AB_11212161 | Chicken | Polyclonal | 1:1000 (72 h, RT) |
| MAP-2 | Proteintech | 17490-1-AP | AB_2137880 | Rabbit | Polyclonal | 1:1000 (72 h, RT) |

**Supplementary Table 1.** Primary antibodies used in this study and immunohistochemical (IHC) application data.

#### Animals

Mice were injected unilaterally into the striatum, with the exception of mice used for preparing electron microscopy samples, which were injected bilaterally. Injections were performed using 3-month-old wild-type C57BL/6J mice (order number #000664, RRID:IMSR_JAX:000664) obtained from Jackson Laboratories (Bar Harbor, ME, USA) and 5-8 weeks old $Snca^{GFP}$ knock-in mice (order number #035412 JAX 035412, RRID:IMSR_JAX:035412).[1] The Snca-GFP knock-in mouse line was used to confirm the presence of pathology for electrophysiological recordings. Both female and male mice were included in the study unless otherwise stated. All animal procedures undertaken in this study to generate SEQUIN, immunochemistry, electron microscopy, and electrophysiology data were approved by the respective University Institutional Animal Care and Use Committees at Yale University, the University of Alabama at Birmingham, and the Washington University School of Medicine and complied with the National Institutes of Health guidelines for the Care and Use of Laboratory Animals.

#### Preparation of recombinant α-synuclein monomer and preformed fibrils for stereotaxic injection

Monomeric mouse α-synuclein was expressed in *E. coli* and purified, followed by endotoxin removal using Pierce LAL high-capacity endotoxin removal resin to achieve levels below 0.1 EU/µg, as previously described.[2] The monomer was stored at −80°C. Preformed fibrils (PFFs) at a concentration of 5.0 mg/ml were produced from monomeric α-syn by agitation in 150 mM KCl/50 mM Tris-HCl at 37°C for seven days.[3,4] PFFs in a volume of 22-25 µl were placed in a 1.5 ml sonicator tube (Active Motif, NC0869649) and sonicated in a Qsonica Q700 sonicator (Newtown, CT) using a cuphorn for 15 min at an amplitude of 30, with a pulse on and off for 3 sec and 2 sec, respectively. The required size range of resulting PFF fragments from 20 to 70 nm was verified by dynamic light scattering on a DynaPro NanoStar Dynamic Light Scattering Detector (Wyatt Technology). PFF samples were stored at -80°C until needed. On the day of injection, the sonicated PFFs were thawed at room temperature and re-sonicated in a Qsonica Q700 sonicator for 15 min at an amplitude of 40, with a pulse on and off for 3 s, respectively, at 10°C and used within 8 h to avoid self-aggregation over time. The PFFs were kept at room temperature during the injections. In contrast, the monomer was kept on ice during the series of injections into multiple animals to prevent fibril aggregation and spun at 20,000 *g* at 4°C with the supernatant used for injections.

#### Stereotaxic injections of α-synuclein monomer and preformed fibrils for SEQUIN and Electron Microscopy analyses

C57BL/6J mice were deeply anesthetized using an isoflurane vaporizer (Harvard Apparatus Mobile Anaesthesia System, model 750239) and placed on a stereotactic frame (Kopf, model 900 LS). Mice were injected with either α-syn monomer or PFFs into the right forebrain at coordinates +0.2 mm relative to Bregma, +2.0 mm from midline, and -2.6 mm beneath the dura to target the dorsal striatum. Injected amounts were 2.0 µl at 5 µg/µl (Nanoject III microinjector, Drummond Scientific; rate, 20 nl/s over 100 cycles with a delay of 8 s between injection pulses). We previously showed that monomer injections did not produce α-syn pathology[3] and did not alter overall structure and abundance of synapses[5] compared to vehicle-injected mice. Thus, PBS injection was used as a control to study and analyse ultrastructural changes by electron microscopy. The needle remained undisturbed for 10 min after protein delivery into the dorsal striatum and was then gradually withdrawn to prevent backflow or leakage into the cortex. The skull was closed with sutures, and the mice were placed on a heating pad to recover until they regained consciousness, then returned to their home cages.

#### Stereotaxic injections of α-synuclein monomer and preformed fibrils for electrophysiological analyses

For slice electrophysiology, 5-8 week old $Snca^{GFP}$ knock-in mice were anesthetized with isoflurane in a chamber, mounted in a stereotaxic frame, and continually maintained under isoflurane anaesthesia using a vaporizer (Harvard Apparatus). Anesthetized mice were unilaterally injected with either monomer or PFF in the right dorsal striatum using the coordinates 0.2 mm posterior to Bregma, 2.0 mm lateral to the midline, and at a depth of 2.6 mm from the dura. Injected amounts of either α-syn monomer or PFFs for electrophysiological analyses were 1.6 µl at 5 µg/µl (Higley laboratory). Injections were performed using a Nanoject III microinjector (Drummond Scientific) at a rate of 20 nl/s over 80 cycles with a delay of 7 seconds between cycles. The pipette was left in place for 10 minutes after the injection to prevent backflow while withdrawing. The incisions were sealed with sutures, and the mice recovered on a heating pad until they were conscious. The mice were then returned to the colony and used for experiments for 5-6 weeks post-injection.

### Stereotaxic co-injection of retrograde AAV-GFP and α-synuclein preformed fibrils for axonal labelling

To retrogradely label axons and neurons projecting to the injection site and simultaneously induce α-Synuclein pathology, pAAV-CAG-GFP (Addgene viral prep # 37825-AAVrg; RRID: Addgene_37825) was co-injected with PFFs. The virus was added to PFFs after sonication to a final titer of $8 \times 10^8$ GC/µl. A volume of 2.5 µl was injected into the dorsal striatum (5 µg total PFF; $2 \times 10^9$ GC AAVrg). Brains were collected 2 weeks after injection, sectioned at 40 µm, and immunostained for p-α-synuclein to detect axonal pathology.

### Tissue processing for synapse quantification

Synaptic loci of the prefrontal cortex were analysed in layer V of the secondary motor cortex M2. We processed tissue for SEQUIN imaging as described, with modifications.[6,7] In brief, mice injected with either monomer or PFFs were analysed at 1, 3, or 6 months post-injection. Mice were deeply anesthetized by intraperitoneal ketamine (100 mg/kg) and xylazine (10 mg/kg) cocktail injection and transcardially perfused with ice-cold 1x Phosphate Buffered Saline (PBS), followed by ice-cold 4% formaldehyde (PFA) in PBS (pH 7.6). The brains were then extracted and post-fixed for 24 h at 4°C in 4% PFA in PBS and stored in PBS at 4°C. Fixed brains were sectioned coronally into 40 µm thick slices using a vibratome (Leica VT1000 S) or 50 µm thick slices using a microtome (Microm HM430). Brain slices were stored in 24-well plates in cryoprotectant buffer (PBS containing 30% ethylene glycol, 30% sucrose, and 0.02% sodium azide) at −20°C.

Slices with prefrontal cortex were washed 3 times for 5 min each in PBS, then blocked in 20% normal goat serum (Jackson ImmunoResearch) in PBS for 1 h at room temperature (RT). Slices were then immunolabeled with primary antibodies, washed 3 times in PBS, followed by Alexa Fluor-labelled secondary antibodies prepared in PBS containing 10% NGS and 0.30-0.45% Triton X-100 in the dark (details on antibodies used and conditions are listed in Table 1). Sodium azide (0.02–0.05%) was added during primary and secondary antibody incubation. The slices were washed 3 times in PBS to remove the unbound secondary antibodies, counterstained with DAPI in PBS for 10 min, rinsed 3 times in PBS, transferred to charged microscopic glass slides, dried at RT, and rinsed in distilled water to remove PBS residues. The slides were dried at RT and embedded in mounting media freshly prepared by mixing Tris-MWL 4-88 (Electron Microscopy Sciences #17977-150, Hatfield, PA, USA) with AF300 (Electron Microscopy Sciences #17977-25) in a 9:1 ratio. High-precision No. 1.5H cover glasses with a thickness of 170 µm ± 5 µm were used (Marienfeld # 0117580). The cover glasses used for SEQUIN were stain-free, thoroughly cleaned with 100% ethanol, rinsed with distilled water, and air-dried prior to coverslipping. The slides were cured at RT for 3 days.

### Synaptic imaging and analysis

Images for synaptic analyses were obtained, processed, and analysed as previously described[7] using a Zeiss LSM 900 Airyscan Microscope (Carl Zeiss Microscopy) with 63x (numerical aperture 1.4) oil immersion objectives (6-10 images per animal from 3 slices) and processed in 3D Airyscan. In brief, images were acquired at ≥1.8× magnification with XY pixels of 43 nm. Within the Z-stack, the step size ("Interval") was set to 120 nm, and 35 optical sections were captured, giving a total image thickness of 4.08 µm. Within each experiment and comparable group, all sections were imaged using identical microscope settings, ensuring consistency for within-experiment comparisons. Experiments conducted at different time points were performed separately with different settings. Images were processed in 3D Airyscan mode using a fixed filter strength for each experiment (within 6.5–7.5, chosen close to the automatically determined value). Channel alignment was verified by inspecting images for chromatic aberration, and manual corrections were applied in Zeiss ZEN 2.3 SP1 (black, version 14.06.201) as needed. Pre- and post-synaptic puncta were detected using Imaris 3D visualization software (version 10.2, Bitplane), and individual puncta and synaptic loci where they were aligned were quantified through a custom nearest neighbour analysis in MATLAB version R2023b. The custom MATLAB-based code is available on GitHub: https://github.com/KummerLab/SEQUIN.

To quantify synapse density, the frequency distribution of nearest-neighbour pre- and post-synaptic protein pairs was generated for the top 20% intensity of pre- and post-synaptic puncta. For the VGLUT2/Homer1 synaptic loci analysis, the top 60% most intense pre- and post-synaptic puncta were used due to the lower number of VGLUT2. Most VGLUT2 puncta were nearest neighbours to Homer. The frequency distribution of nearest neighbour pairs, which forms a Gaussian distribution and excludes all random non-synaptic pairings, was used to calculate the synaptic locus density. The mean value calculated at the peak of the Gaussian distribution provided the average pre-post synaptic separation. Synapse densities were analysed using linear mixed-effects models with mouse as the unit of inference in SPSS (v31.0.0.0, IBM Corp.). Repeated measurements across cortical areas were modelled with animal and region as random effects to avoid pseudoreplication and reflect biological replication.

#### Synaptic imaging of endogenous α-synuclein

Cortical sections from mouse controls 3 MPI with monomer, co-stained with VGLUT1/α-synuclein or VGLUT2/α-synuclein were imaged using the synaptic imaging protocol described above with minor modifications. Images were acquired in 3D Airyscan mode (strength 5.5), and chromatic aberration was corrected. Background subtraction was performed in Imaris, and the Coloc function was applied to quantify colocalization of VGLUT1 with α-synuclein and, in a separate analysis, VGLUT2 with α-synuclein. Threshold parameters were determined independently for each channel using the automatic thresholding function (calculate threshold). Colocalization was then assessed using the Build Coloc Channel function in Imaris, and parameters including Manders' coefficient and the percentage of VGLUT1- or VGLUT2-positive voxels above threshold colocalized with α-synuclein were determined. Statistical analysis was performed using a linear mixed-effects model.

#### Synuclein pathology progression analysis

Mouse cortical slices obtained at 1, 3, and 6 months post-injection of PFF into the striatum were co-stained for DAPI and phosphorylated α-synuclein (pS129 α-syn) to measure synuclein pathology. Ipsilateral cortical sections, i.e., from the same side where PFF was injected into the striatum, were used to analyse phospho-α-synuclein pathology. Tiled cortical images were acquired using a spinning disk Dragonfly microscope (Andor–Oxford Instruments) equipped with a Zyla CMOS camera and a 20× air objective. The stitched 3D images with z-stacks of 10 µm were analysed using IMARIS 3D visualization software (v10.2, Oxford Instruments). The surface module was used to quantify the intensity and size of phospho-α-synuclein aggregates (surface detail, absolute intensity, and threshold), as previously described for the volumetric quantification of synuclein pathology.[8] Synuclein pathology progression, measured as changes in aggregate intensity and volume across time points, was analysed using ordinary one-way ANOVA followed by Holm–Šidák post hoc test for multiple comparisons.

#### Localization of α-synuclein aggregates in axons and dendrites

To identify whether neuritic phospho-α-synuclein aggregates were localized to axons or dendrites, cortical tissue sections were co-labelled with antibodies targeting pS129 α-syn, NF-H to label axons, and MAP-2 to label dendrites. Sections were mounted and coverslipped using MWL 4-88 and 1.5H coverslips for super-resolution imaging using a 63X 1.4NA objective and Zeiss LSM 980 with Airyscan detector. 16 13.8 µm z-stacks were acquired from each animal, covering 159.2µm X 159.2µm. To perform colocalization analysis, four binary masks were generated from the raw imaging data: neuronal soma, neuritic α-syn, axons, dendrites. Given the expected heterogeneity in morphology and labelling across the structures being colocalized, masking parameters were optimized for each type of structure individually. Neuronal soma were identified using the MAP-2 images, which were downsampled 25%, converted to 8-bit, Gaussian smoothed, and then processed with Cellpose v4.1.0 using parameters set to: diameter:150, flow threshold:3, cellprob threshold:0, lower:1, upper:99, niter dynamics:2000, stitch threshold:1, flow3D smooth:0, anisotropy:1, min size:500. Phospho-α-synuclein aggregates were masked based upon intensity thresholding. NF-H labelled axons were masked by first using a Gaussian 3D local threshold with a window size of 3x35x35 (zyx), sigma of 3x9x9 (zyx), and a constant intensity cutoff of –0.75. After this initial masking step, the mask was refined by removing pixels within the mask that were below the $5^{th}$ percentile. MAP-2 labelled dendrites were masked by first setting an intensity threshold set to the $75^{th}$ percentile to create a coarse mask, and then refining this mask by removing pixels within the mask that were below the $5^{th}$ percentile. All masks were compared to the original images to confirm the accuracy. Pixels that were identified by both the axon and dendrite masks were removed from analysis, given ambiguity in their identity. This removal step was confirmed to not significantly alter the number of identified axons or dendrites, nor the magnitude or interpretation of effects,

For every identified phospho-α-synuclein aggregate, the area of coverage by both the axon and dendrite masks was calculated. Aggregates were classified as being either axon or dendrite associated when the area coverage exceeded 50% of the other classification type. To confirm the results of subtype classification were based under the underlying structural biology of the tissue, the analysis was repeated under conditions, which broke any spatial relationship between the labelled aggregates and axon/dendrite masks. This was performed by rotating the axon and dendrite masks 90 degrees relative to the α-syn mask three separate times in the XY plane and repeating the colocalization analysis. The masks were also flipped in Z and the four rotational positions were also processed. This yielded 7 different iterations for each tile where the spatial relationship of α-syn, relative to the axon and dendrite masks, was known to have no biological meaning.

#### Tissue processing for cortical mesoscans

Tissue staining for cortical mesoscans was performed using SEQUIN modifications.[6,7] with minor modifications. Prior to the application of primary antibodies, sections were rinsed with phosphate-buffered saline (PBS) and then blocked with 20% normal goat serum (Vector Labs S-1000) in PBS with 0.3% Triton X-100 (Sigma-Aldrich X100) and 0.05% sodium azide for 1 hour. The blocking solution was then replaced with primary antibodies in 10% normal goat serum in PBS with 0.3%

Triton X-100 (Sigma-Aldrich X100) and 0.05% sodium azide for 3 days. To reduce the possibility of off-target labelling when performing mouse-on-mouse staining, the antibody directed against phosphorylated α-syn was preconjugated to Alexa-647 using Thermo Alexa Fluor™ 647 NHS Ester (succinimidyl ester-conjugated) (Thermo cat# A20006) and purified using Zeba spin desalting columns (Thermo Cat# 89883). Following primary incubation, tissue sections were thoroughly rinsed, and then the secondary antibodies Goat anti-Rabbit 568 (Thermo cat# A-11011) and Donkey anti-Guinea Pig 488 (Jackson ImmunoResearch Cat# 706-545-148) were applied and incubated overnight. The following day, sections were thoroughly rinsed, and the nuclear counterstain DAPI was applied. Sections were rinsed again and then mounted on charged slides and coverslipped with Mowiol 4-88 + AF300 (9:1 ratio, Electron Microscopy Sciences, #17977-150 and #17977-25) and 1.5h coverslip (Marienfeld #0107242).

#### Acquisition of cortical mesoscans and image analysis

Images for analyses of synapses on population levels were acquired using a Zeiss LSM 980 microscope using the 8Y multiplex scanning mode, allowing for increased scanning speed while maintaining sufficient super-resolution imaging capabilities to perform synaptic localization analysis. Regions of interest were outlined in Zen Blue, and a 3x3 array of focus points were generated. At each focus point, the centre of the z-stack was manually set to the centre of the tissue section, and a horizontal plane focus map was created. Every section was imaged with identical microscope imaging parameters. During each imaging session, images from a 'Channel Alignment' section were acquired for the measurement and correction of chromatic aberration.[7] Following the acquisition, images were Airyscan processed using 3D mode with a strength of 4.0.

Image analysis was performed using https://github.com/KummerLab/SEQUIN. Since the SEQUIN-based selection of loci is intensity-based and allows for direct within-animal comparisons across regions, for each tile scan, the 5$^{th}$ percentile of intensity for each synaptic channel was calculated prior to spot detection and then applied to all tiles. Once the synaptic loci had been identified, all subsequent metadata was obtained from the raw imaging data. To improve the identification and separation of synaptic and non-synaptic pairs during SEQUIN analysis, a gradient-boosting machine learning classifier was trained using LightGBM[9] available at https://github.com/microsoft/LightGBM, and metadata on the colocalized pre- and post-synaptic loci from original SEQUIN based methods. Direct comparison of the machine learning classifier versus previous SEQUIN methods revealed matching results for standard intensity cutoff ranges. Notably, the classifier was able to separate the primary synapse peaks from the entire dataset even as dimmer puncta were included in the analysis, similar to two-Gaussian deconvolution, while also maintaining the identity of each punctum which is required for all subsequent analyses in these studies.

Quantification of synapse-associated p-α-syn was performed on synaptic loci identified using SEQUIN. All images received a fixed cutoff background subtraction of the p-α-syn labelling, calculated from monomer-treated animals, which do not exhibit prominent p-α-syn aggregation and are thus a measure for non-specific labelling. For each synaptic pair of pre- and post-synaptic puncta, a 200 nm field was defined around the centroid and the intensity of the p-α-syn channel was measured. The parenchymal intensity of p-α-syn was quantified using the intensity of p-α-syn outside of Lewy neurites but within the same image field. For the comparison of synaptic and parenchymal p-α-syn levels in PFF-treated animals, given the regionally specific aggregation patterns of p-α-syn, the top 20% of tiles by p-α-syn level were identified for analysis. These tiles were manually verified to represent the areas with prominent p-α-syn aggregation. Statistical analysis of mesoscan data involving multiple cortical areas was performed using a linear mixed-effects model (fitlme) in MATLAB version R2023b. This approach accounted for the requirement for repeated imaging from the same animal and allowed for animal and region of interest to be specified as random effects during the analysis. Heatmaps of synaptic density and p-α-syn levels were generated for each animal and registered to a common reference section using a landmark-based affine registration (ImageJ, NIH). Registered heatmaps for each group were combined to generate final group-wise heatmaps.

For quantification of synapse-associated p-α-syn as a function of distance to Lewy neurites, p-α-syn-positive neurites were first masked. Neurite masks were generated by using a size exclusion difference-of-gaussian filter followed by an intensity cutoff equivalent to the $95^{th}$ percentile of p-α-syn intensity for areas of high pathology, allowing for uniform detection of neurites across areas of differing magnitudes of pathology and treatments. For each synaptic centroid, the nearest voxel within a masked Lewy neurite was then measured. For each masked Lewy neurite, the intensity of p-α-syn within the neurite mask was measured, then the mask was dilated 800 µm, and the density of synaptic loci within the dilated area was quantified. The dilation radius of 800 µm corresponds to the average neurite-synapse separation distance and is the separation distance where increased synaptic p-α-syn begins to be detected.

Quantification of synaptic alterations within cortical columns was performed using a combination of atlas registration for identification of cortical layers using QuickNII and VisuAlign available at https://scicrunch.org/resolver/RRID:SCR_017978, followed by a semi-automated segmentation of cortical columns. Layer-specific alterations in synaptic density were analysed with a Mann-Whitney test on layer-specific sub-regions of individual imaging tiles. The boundaries between layers I vs II/III and layers II/III vs V of the entire secondary motor cortex M2 were traced manually and five cortical columns per animal were identified. Synaptic p-α-syn was measured in 200 nm fields around identified synaptic centroids and total p-α-syn was measured from maximum intensity projections of the same cortical mesoscans used for synaptic analysis. Given that cortical column and layer specific ROI boundaries could subdivide individual z-stack tiles, only tiles that maintained at least half of their image were kept. To ensure correlations were performed on sufficient imaging data, only cortical columns that contained at least two image tiles were included in the analysis.

### Electron microscopy and image analysis

Electron microscopy and image analysis were conducted as described previously. Briefly, male mice (n=four PBS, n=four PFF) were anesthetized using isoflurane and perfused with phosphate-buffered saline (PBS) followed by 2.5% glutaraldehyde and 2% paraformaldehyde (pH 7.5). Brains were fixed in 2.5% glutaraldehyde and 2.0% paraformaldehyde in PBS for 2 hours at room temperature and then transferred to 4°C for 16 h. Embedding was performed using 2% 255-bloom calf skin gelatin with 3% agarose in PBS. A vibratome was used to cut 200 µm sections followed by dissection of M2 cortex, and immersion in 2% paraformaldehyde in 0.1 M cacodylate buffer at pH 7.4. The samples were shipped in 0.1 M phosphate buffer to the Centre for Cellular and Molecular Imaging EM Core facility at Yale Medical School.

Post-fixation was performed using 1% OsO4 and 0.8% potassium ferricyanide in 0.1 M cacodylate followed by en bloc staining with 2% aqueous uranyl acetate, dehydration in a graded series of ethanol up to 100%, substituted with propylene oxide, and embedded in EMbed 812 resin (Electron Microscopy Sciences, Hartfield, PA). Following polymerization of sample blocks at 60°C, 60 nm sections were cut using a Leica ultramicrotome (UC7) and post-stained with 2% uranyl acetate and lead citrate. Imaging was performed with FEI Tecnai transmission electron microscope at an 80 kV accelerating voltage, and digital images were acquired with an AMT NanoSprint15 MK2 camera (Advanced Microscopy Techniques, Woburn, MA).

Asymmetrical synapses were quantified by a researcher blinded to experimental conditions. Synapses were excluded from analysis if they exhibited a poorly defined postsynaptic density (PSD), if the PSD signal was so intense that it obscured both membranes, if the presynaptic terminal contained fewer than 4 synaptic vesicles, or if synapses were at the edge of the frame. A convolution neural network trained on mouse synapses was utilized to measure the area of individual synaptic vesicles.[10] For the measurement of synaptic cleft width, the two edges of the PSD adjacent to the presynaptic terminal were determined, and the distance between the PSD and presynaptic membrane was measured at the midpoint between these two edges. The PSD was manually traced using ImageJ, and the length was measured using the segmented line tool.

For statistical analyses, the 5% minimum and 5% maxima were trimmed because histograms revealed right-skewed data, and a Box-Cox transformation was applied. Data were analysed using a linear mixed model with mouse number as "subjects" and synapse number as repeated measures; compound symmetry variance structure was used with treatment as a fixed effect.

### Electrophysiology

The Snca-GFP knock-in mouse line[1] was used for recordings to confirm the presence of pathology and locate density burden in the cortical area where recordings were obtained. Predominantly neuritic Snca-GFP aggregates were observed at the time point used for recordings. Mice striatally injected with monomer or PFFs as described above were at 5-6 weeks post-injection deeply anesthetized with isoflurane and transcardially perfused with oxygenated (95% $O_2$/5% $CO_2$) ice-cold choline artificial cerebrospinal fluid (Choline ACSF) containing (in mM): 110 choline, 25 $NaHCO_3$, 1.25 $NaH_2PO_4$, 2.5 KCl, 7 $MgCl_2$, 0.5 $CaCl_2$, 20 glucose, 11.6 sodium ascorbate, 3.1 sodium pyruvate. The perfused mice were decapitated, and the brains were quickly removed for acute slice (300 μm thick) preparation from the injected (PFF or monomer) hemisphere. Acute slices were prepared in the coronal plane from the region neighbouring to the injection site, and the recordings were conducted from the primary somatosensory cortex. Slices were incubated in warm ACSF (32°C) containing (in mM): 127 NaCl, 25 $NaHCO_3$, 1.25 $NaH_2PO_4$, 2.5 KCl, 1 $MgCl_2$, 2 $CaCl_2$, and 20 glucose bubbled with 95% $O_2$/5% $CO_2$. After incubating at 32°C for 30 minutes, the slices were stored at room temperature (RT). Whole-cell recordings were carried out at RT from layer II/III pyramidal cells located in proximity to GFP-labelled aggregates. Pyramidal cells were identified under differential interference contrast by their pyramidal somatic shape and prominent apical dendrite. Series resistance was maintained at less than 25MΩ and uncompensated. Recordings were made using a Multiclamp 700B amplifier (Molecular Devices), filtered at 2 kHz, and digitized at 10 kHz using custom software.[11] To record miniature postsynaptic currents, 1 μM tetrodotoxin (TTX) was added to the bath. Glass electrodes (3.0-3.5 MΩ) were backfilled with an internal solution containing (in mM): 126 cesium gluconate, 10 HEPES, 10 sodium phosphocreatine, 4 $MgCl_2$, 4 $Na_2ATP$, 0.4 $Na_2GTP$, 1 EGTA (pH 7.3 with cesium hydroxide). Cells were voltage-clamped at -70mV to record miniature excitatory postsynaptic currents (mEPSCs) and at 0mV to record miniature inhibitory postsynaptic currents (mIPSCs).

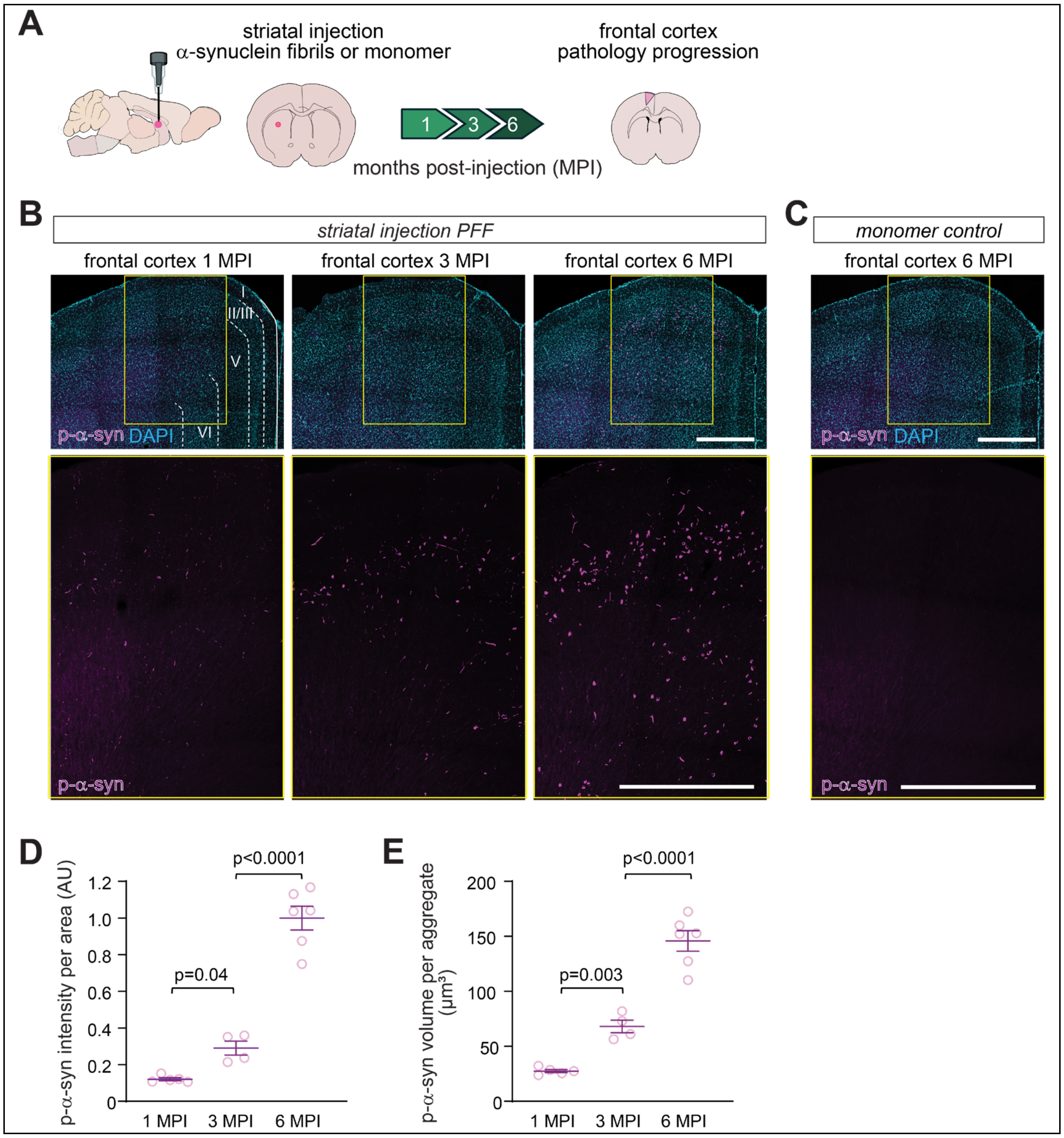


**Supplementary Fig. 1. Progressive development of cortical α-synuclein pathology following PFF injection in mice.**

(**A**) Diagram of striatal injection of α-syn pre-formed fibrils (PFFs) in mice, resulting in progressive pathology in cortical neurons projecting to the striatum. Injection of α-syn monomer served as control.

(**B**) Representative images of α-syn aggregates detected by staining of frontal cortex for p-α-syn at 1, 3, and 6 months post-injection (MPI) of PFF into the striatum. Top, co-staining of p-α-syn (magenta) and DAPI (cyan). Bottom, p-α-syn signal alone for the boxed region shown in the merged DAPI/p-α-syn image above. Cortical α-syn aggregates were apparent at 1 MPI as measured by immunostaining for p-α-syn. Scale bars, 500 μm.

(**C**) No p-α-syn pathology was detected 6 months post-injection of monomer. Scale bars, 500 μm.

(**D, E**) Quantification of p-α-syn staining per frontal cortex area from images as in (**B**) showed pathology progression from 1 MPI, with the intensity (**D**) and mean volume (**E**) of p-α-syn

aggregates increasing at 3 and 6 MPI. (**D** and **E**, n=four–six animals per group, with two–three slices per animal)
Results are presented as mean ± SEM and analysed by ordinary one-way ANOVA with multiple comparisons test and considered statistically significant with $p<0.05$.

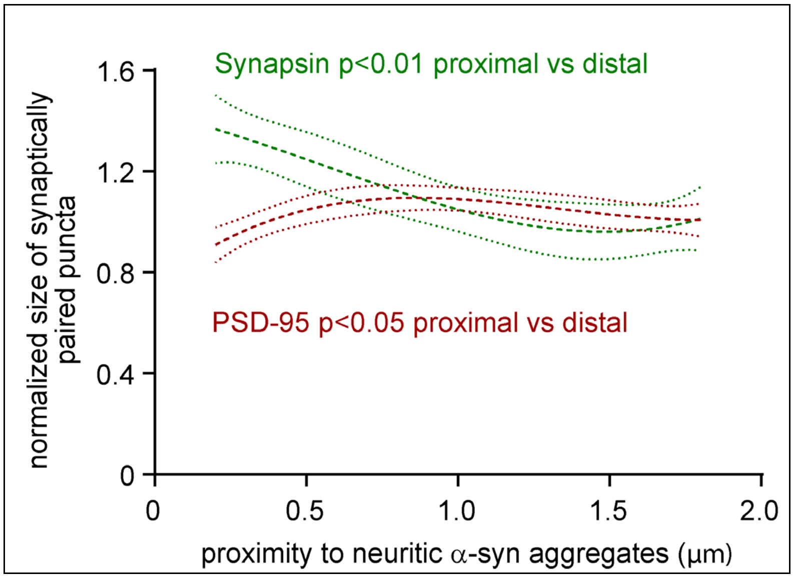


**Supplementary Fig. 2. Synaptic puncta in close proximity to α-synuclein aggregates exhibit changes in size.**

Synaptic loci, imaged by Airyscan super resolution microscopy and identified with SEQUIN, were analysed by imaging tile and grouped based upon the distance to the nearest p-α-syn aggregate. Fig. 2 shows representative images. Comparing synaptic loci in close proximity to a p-α-syn aggregate (within 0.4 μm) to those distal (1.6-2.0 μm) showed a significant enlargement of synapsin labelled pre-synaptic terminals, while the size of the PSD-95 labelled post-synaptic specializations was reduced. Statistical analyses were performed using a linear mixed-effects model. ($p=0.003$ proximal vs distal synapsin puncta, $p=0.031$ proximal vs distal PSD-95 puncta, $n=863$ image tiles from three mice)

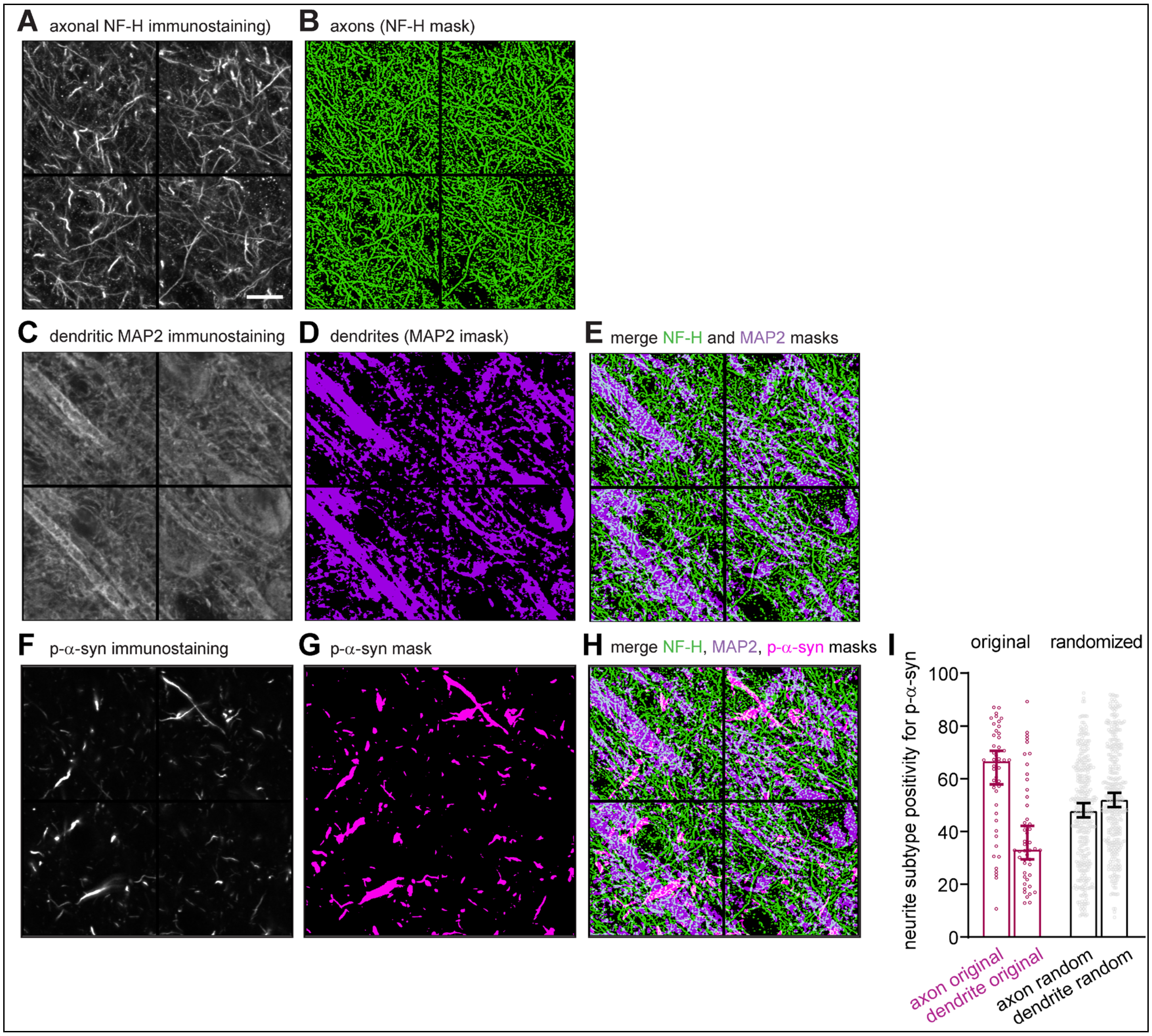


**Supplementary Fig. 3. Representative images and corresponding masks used for the localization of p-α-synuclein aggregates to axons and dendrites.**

Airyscan super-resolution microscopy was performed at 1 MPI of PFFs in M2 cortex to colocalize axons, dendrites, and p-α-syn aggregates.

(**A**) Four representative areas of a tilescan showing maximum intensity projection of the axonal marker Neurofilament-H (NF-H). Scale bar in (**A**), 10 µm; also applies to (**B-H**).

(**B**) Binary mask of the signal from NF-H positive axon generated from the image shown in (**A**).

(**C**) Maximum intensity projection of the dendritic marker MAP-2 in the area shown in (**A**).

(**D**) Binary mask of the signal from MAP2-positive dendrites generated from the image in (**C**).

**(E)** Merged image of the binary masks in (**B**) and (**C**) shows separation of NF-H positive axons and MAP2-positive dendrites.

**(F)** Maximum intensity projection of p-α-syn aggregates in the area shown in (**A**).

**(G)** Binary mask of the signal of p-α-syn aggregates generated from the image in (**E**).

**(H)** Merged image of all binary masks in (**B**), (**C**), and (**E**) for quantification of the extent of localization of p-α-synuclein aggregates to NF-H positive axons and MAP2-positive dendrites.
(**I**) Quantification of axonal and dendritic localization of p-α-synuclein in contralateral cortex reveals more frequent detections in axons compared to dendrites, and fewer overall colocalizations compared to ipsilateral cortex. Statistical analysis was performed on 16 image tiles per animal from three mice using a linear mixed-effects model. Randomization analysis was also performed using multiple rotations of the axonal and dendritic mask images relative to the p-α-synuclein image. This demonstrated that the higher frequency of axonal versus dendritic p-α-syn colocalizations did not arise as a result of unrelated features, such as total coverage of the axonal or dendritic masks.

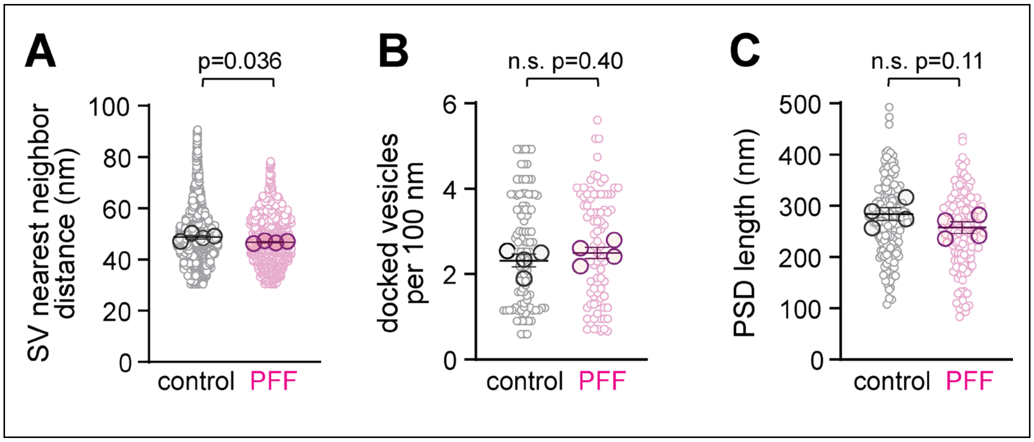


**Supplementary Fig. 4. Ultrastructural analysis of synaptic pathology.**
(**A**) Analysis of EM images determined a modest but significant reduction in the nearest neighbour distances of synaptic vesicles in mice injected with PFF compared to controls. (PBS, n=8810 synapses; PFF, n=7576 from n=four mice per group).
(**B**) The number of docked vesicles along the presynaptic membrane apposed to the PSD was unchanged between the two groups. (PBS, n=334 synapses; PFF, n=309 from four mice per group)
(**C**) No significant change in the length of the PSD was observed. (PBS, n=498 synapses; PFF, n=453 from four mice per group).

Results are presented as mean ± SEM, calculated from mouse means. Small circles represent individual image measurements, whereas large circles indicate the mean per mouse. Data were analysed after Box-Cox transformation using linear mixed models with synapses nested within each mouse, as in Fig. 4E and F.

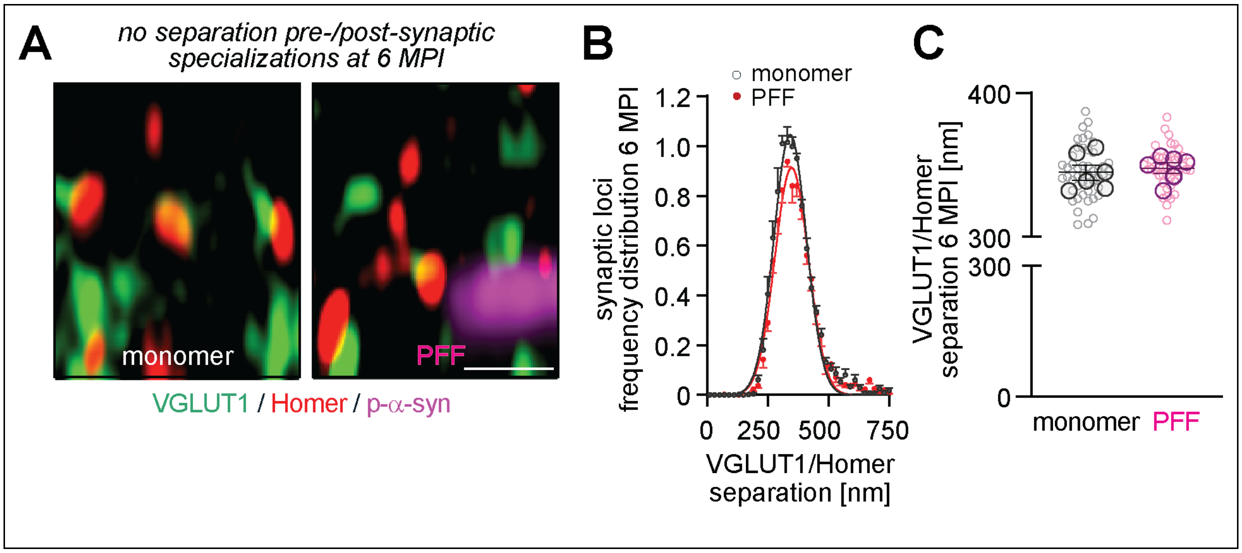


**Supplementary Fig. 5. No evidence for structural aberrations at VGLUT1/Homer synapses in the presence of cortical α-synuclein aggregates at later stages of pathology.**

(**A**) Representative high-resolution Airyscan images from the frontal cortex M2 layer V. Images were acquired 6 MPI of monomer control (left) or PFF (right) and show paired pre- and post-synaptic excitatory VGLUT1 (green) and Homer (red) specializations, with p-α-syn in magenta. Scale bar, 1 µm.

(**B, C**) No difference in spatial separation of VGLUT1/Homer in PFF- compared to monomer control-injected mice at 6 MPI. Nearest-neighbour distances were determined by analysing the Gaussian distribution (**B**) and mean separation distances (**C**) of synaptically paired puncta using SEQUIN method. The Gaussian distribution of synaptic loci was normalized to set the amplitude to 1 for monomer-injected control mice. The graph shows separation distances for individual synaptic pairs after nearest-neighbour analysis.

Results are presented as mean ± SEM, where SEM is calculated from mouse means. Small circles in (**C**) represent individual image measurements; large circles indicate the mean per mouse. Group differences were assessed using a linear mixed-effects model, with group as a fixed effect and mouse as the unit of inference. Statistical significance was defined as $p<0.05$. (n=six animals/group, six–eight slices per animal)

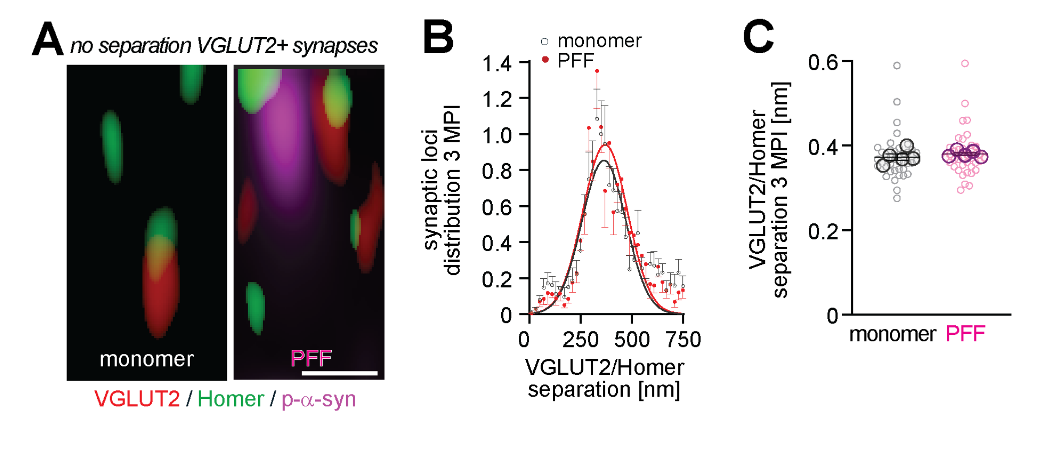


**Supplementary Fig. 6. α-Synuclein pathology does not impact VGLUT2-positive long-range inputs.**
(**A**) Representative high-resolution images from the cortex at 3 MPI showing paired pre- and post-synaptic VGLUT2 (green) and Homer (red). p-α-syn is shown in magenta. Tissues were obtained after injection of monomer as control (left) and PFF (right). Scale bar, 1 µm.
(**B, C**) Separation distances of paired VGLUT2/Homer specializations were unchanged 3 MPI of PFF compared to monomer control. Nearest-neighbour separation distances between pre- and postsynaptic proteins (**B**) were determined by analysing the Gaussian distribution of VGLUT2 and Homer paired synaptic loci signals using SEQUIN. The Gaussian distribution of synaptic loci was normalized to set the amplitude to 1 for monomer-injected control mice. Mean separation distances (**C**) for individual synaptic VGLUT2/Homer pairs are not distinguishable after nearest-neighbour analysis for monomer control vs. PFF injection. (n=five animals/group, six-eight slices per animal).

Results are presented as mean ± SEM, calculated from mouse means. Small circles represent individual image measurements, whereas large circles indicate the mean per mouse. Group differences were assessed using a linear mixed-effects model, with group as a fixed effect and mouse as the unit of inference. Statistical significance was defined as $p < 0.05$.

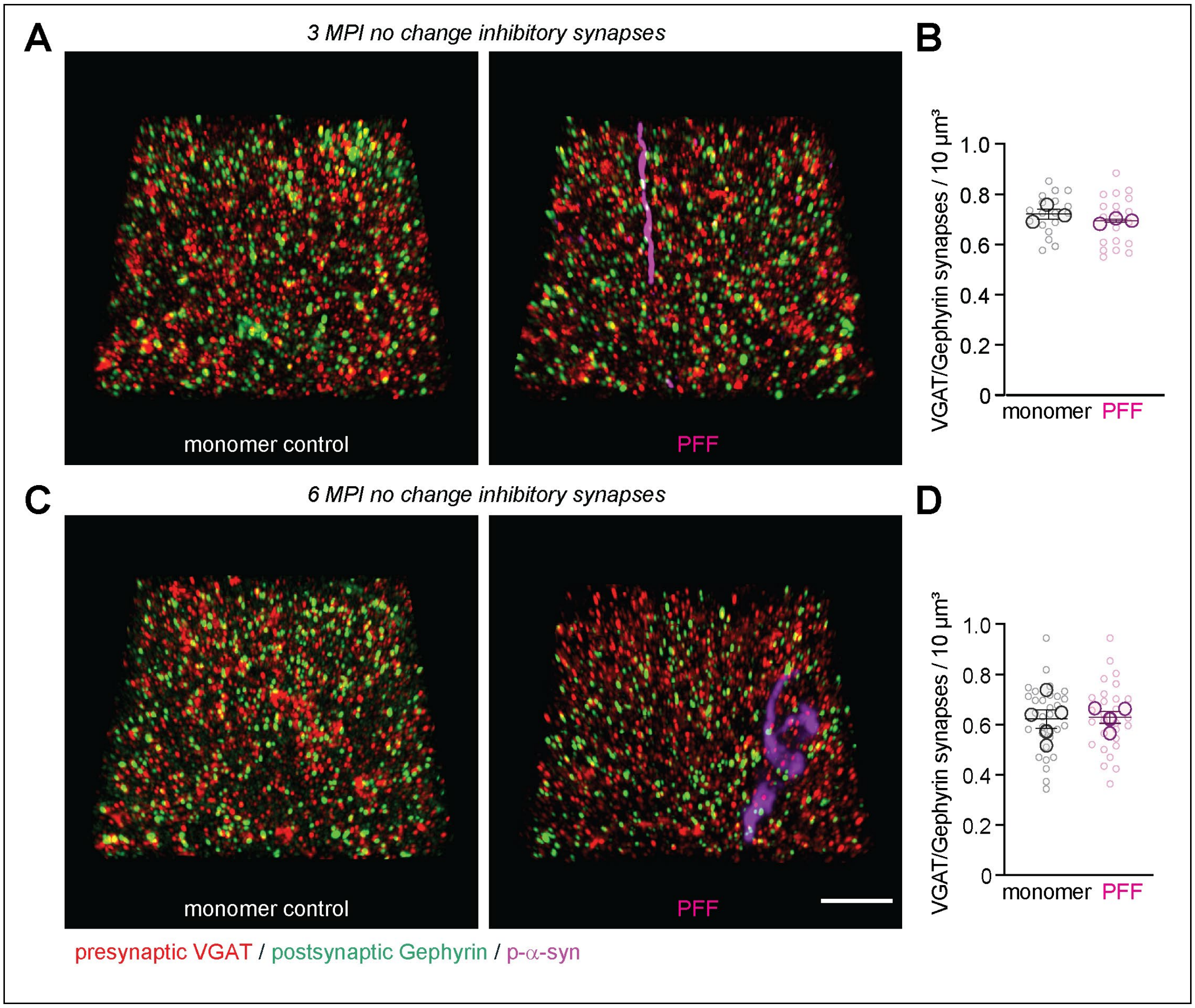


**Supplementary Fig. 7. Inhibitory synapses remain unaffected during α-synucleinopathy progression.**

(**A, C**) Representative 3D high-resolution immunostaining of inhibitory presynaptic VGAT (red) and postsynaptic Gephyrin (green) in frontal cortex M2 at 3 MPI (**A**) and 6 MPI (**C**) after monomer (left) or PFF injection (right), with p-α-syn shown in magenta. Immunostainings were obtained by Airyscan microscopy. Scale bar, 5 µm.

(**B**) Inhibitory synapse density determined by measuring VGAT and Gephyrin puncta paired into synapses was unchanged 3 MPI PFF compared to monomer. Synaptic density was calculated from images as in (**A**) using SEQUIN by nearest-neighbour analyses of puncta. (n=three animals/group, six-seven slices per animal)

(**D**) No change in inhibitory synapse density was observed 6 MPI monomer compared to PFF after analysis of images as in (**C**). (n=four-five animals/group, six-seven slices per animal)

Results are presented as mean ± SEM, calculated from mouse means. Small circles represent individual image measurements, whereas large circles indicate the mean per mouse. Group differences were assessed using a linear mixed-effects model, with group as a fixed effect and mouse as the unit of inference. Statistical significance was defined as $p<0.05$.

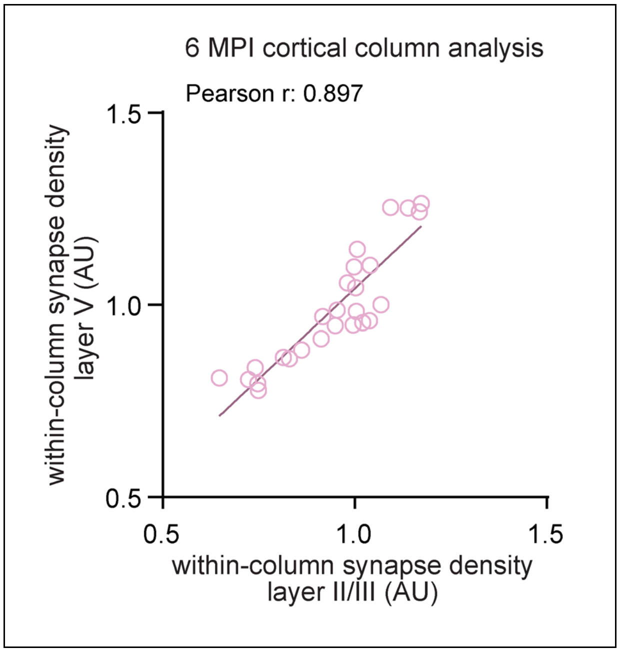


**Supplementary Fig. 8. Excitatory synaptic density in PFF-injected mice is correlated across cortical layers at 6 MPI.**

Synaptic density in layers II/III and V at 6 months after PFF injection exhibited a strong correlation within cortical columns. Analysis was performed as in Figure 7F. (n=4-5 cortical columns per animal from 6 animals; Pearson correlation coefficient provided in the panel)
